# Three-dimensional synaptic organization of layer III of the human temporal neocortex

**DOI:** 10.1101/2021.01.18.427131

**Authors:** Nicolás Cano-Astorga, Javier DeFelipe, Lidia Alonso-Nanclares

**Author notes:** Correspondence to: L. Alonso-Nanclares.

## Abstract

In the present study, we have used Focused Ion Beam/Scanning Electron Microscopy (FIB/SEM) to perform a study of the synaptic organization of layer III of Brodmann’s area 21 in human tissue samples obtained from autopsies and biopsies. We analyzed the synaptic density, 3D spatial distribution, and type (asymmetric/symmetric), as well as the size and shape of each synaptic junction of 4945 synapses that were fully reconstructed in 3D. Significant differences in the mean synaptic density between autopsy and biopsy samples were found (0.49 and 0.66 synapses/μm^3^, respectively). However, in both types of samples (autopsy and biopsy), the asymmetric:symmetric ratio was similar (93:7) and most asymmetric synapses were established on dendritic spines (75%), while most symmetric synapses were established on dendritic shafts (85%). We also compared several electron microscopy methods and analysis tools to estimate the synaptic density in the same brain tissue. We have shown that FIB/SEM is much more reliable and robust than the majority of the other commonly used EM techniques. The present work constitutes a detailed description of the synaptic organization of cortical layer III. Further studies on the rest of the cortical layers are necessary to better understand the functional organization of this temporal cortical region.

## Introduction

Several electron microscope (EM) studies have been performed to estimate the density and types of synapses in the human temporal cortex with a range of different standard EM techniques (Davies et al. 1987; Huttenlocher and Dabholkar 1997; Tang et al. 2001; Alonso-Nanclares et al. 2008; Yakoubi et al. 2018, 2019). However, these estimations are usually applied to a relatively small number of EM sections, which increases statistical variability and affects the reliability of the results (reviewed in Merchán-Pérez et al. 2009). Furthermore, in these studies, many synapses cannot be properly classified as asymmetric or symmetric, and they are tagged as “uncharacterized” (DeFelipe et al. 1999). This missing data is critical at the circuit level, since higher or lower proportions of asymmetric and symmetric synapses are linked to differences in the excitatory/inhibitory balance of the cortical circuits (for reviews, see DeFelipe 2015; Froemke 2015; Zhou and Yu 2018; Sohal and Rubenstein 2019).

EM with serial section reconstruction is the gold standard method for studying the synaptic organization. However, it is rather time-consuming and challenging to obtain long series of sections. This is why reconstruction of large tissue volumes is usually not feasible. The introduction of automated or semi-automated EM techniques at the turn of this century represents a major advance in the study of synaptic organization, as long series of consecutive sections can now be obtained with little user intervention (e.g., (Denk and Horstmann 2004; Knott et al. 2008; Smith 2008; Merchan-Pérez et al. 2009; Kleinfeld et al. 2011; Helmstaedter et al. 2013; Kasthuri et al. 2015; Kubota et al. 2018). Regarding the way in which human brain tissue is acquired, there are two main sources: (i) postoperative brain tissue (biopsy) obtained from patients suffering pharmacoresistant temporal lobe epilepsy or from patients with brain tumors; and (ii) from the autopsy of individuals with no recorded neurological or psychiatric alterations.

The advantage of using biopsies is that the resected tissue can be immediately immersed in the fixative. This is why the quality of the immunocytochemical staining at both the light and electron microscopy levels in human biopsy material has been shown to be comparable to that obtained in experimental animals (e.g., del Río and DeFelipe 1994) The problem is that, although brain tissue obtained at biopsy may come from part of a region that is considered ‘normal’ from a pathological and physiological point of view, it is obtained from patients who did have an underlying brain-related disorder. Thus, autopsy may be the sole source of strictly normal tissue. The problem is that there can be a long delay between death and tissue collection (often over 5 hours), and the ultrastructure of the post-mortem brain tissue is generally not well preserved, which makes the tissue unsuitable for detailed quantitative analysis. This is one of the main reasons why synaptic circuitry data for the normal human brain is so lacking. However, we have shown that the 3D reconstruction method using Focused Ion Beam/Scanning Electron Microscopy (FIB/SEM) can be applied to study the synaptic organization of the human brain obtained at autopsy in great detail, yielding excellent results (Domínguez-Álvaro et al. 2018, 2019, 2020; Montero-Crespo et al. 2020).

The main objective of this study is to analyze the synaptic organization of the neuropil of layer III of the anterior part of the middle temporal convolution (Brodmann’s area 21; BA21; see Zilles and Amunts 2010) obtained from both biopsies and autopsies. It has been proposed that this cortical layer is critically involved in the emergence of cognitive functions, such as language (Thomson and Lamy 2007; Xu et al. 2016; Cheng et al. 2017) and most physiological studies in the human neocortex are performed in layers II and III (e.g., see Eyal et al. 2018; Gidon et al. 2020, and references therein). Thus, a detailed microanatomical analysis could be useful to better understand this cortical region by integrating morphological and physiological studies. We used FIB/SEM in normal brain tissue obtained at autopsy with a short postmortem delay (less than 4h) and from ‘normal’ tissue obtained at biopsy (from patients with mesial temporal lobe epilepsy, with no fixation delay) to determine the synaptic organization in this particular brain region.

The present study also allows the comparison of the data from autopsy and biopsy using the same technique (FIB/SEM). Moreover, since this layer has been examined in previous studies using standard EM techniques and stereological methods (Alonso-Nanclares et al. 2008), the present study also aimed to compare standard EM techniques with the FIB/SEM to study the synaptic organization. For this purpose, we studied 4945 synapses, which were fully reconstructed in 3D. In particular, we analyzed the synaptic density and 3D spatial distribution, type of synapses (asymmetric/symmetric), as well as the shape and size of each synaptic junction. In addition, it was possible to determine the postsynaptic targets of 2304 axon terminals. Thus, data from the present work constitutes a detailed description of the synaptic organization in layer IIIA of the human BA21, which may contribute to better understanding the synaptic organization of this layer.

## Materials and Methods

### Tissue preparation

Human brain tissue was obtained from 3 autopsies and 5 biopsies (Table 1). The autopsy samples (with short post-mortem delays of less than 4 hours) were obtained from 2 men and 1 woman (whose ages ranged from 45 to 53 years old) with no recorded neurological or psychiatric alterations (supplied by Unidad Asociada Neuromax, Laboratorio de Neuroanatomía Humana, Facultad de Medicina, Universidad de Castilla-La Mancha, Albacete). The sampling procedure was approved by the Institutional Ethical Committee. This human brain tissue has been used in previous EM studies (Domínguez-Álvaro et al. 2018, 2019, 2020; Montero-Crespo et al. 2020). Biopsy samples were obtained during neurosurgery of epileptic patients suffering pharmaco-resistant mesial temporal lobe epilepsy in the Hospital de la Princesa (Madrid). Intraoperative electrocorticographic recordings revealed that the lateral neocortex of all these patients displayed normal activity. That is, no spikes, sharp waves, or slow activities were observed during intraoperative electrocorticography. A portion of the anterior part of the left temporal lobe was removed. In each case, the patient’s consent was obtained in accordance with the Helsinki Declaration, and all protocols were approved by the ethical committee at the Hospital de la Princesa (Madrid). This biopsy tissue has also been used in previous studies (Alonso-Nanclares et al. 2008).

**Table 1.** Clinical information of the analyzed cases. Causes of death are shown in parentheses in the autopsy cases. TLE: temporal lobe epilepsy.

| **Case** | **Sex** | **Age**<br>**(years)** | **Tissue type** | **Postmortem delay (h)** |
| --- | --- | --- | --- | --- |
| **AB1** | Male | 45 | Autopsy<br>(cardiorespiratory arrest) | <1 |
| **AB2** | Female | 53 | Autopsy<br>(pulmonary septic shock) | 4 |
| **AB3** | Male | 53 | Autopsy (bladder<br>metastatic carcinoma) | 3.5 |
| **H38** | Male | 24 | Biopsy TLE | 0 |
| **H18** | Female | 31 | Biopsy TLE | 0 |
| **H60** | Male | 36 | Biopsy TLE | 0 |
| **H47** | Female | 41 | Biopsy TLE | 0 |
| **H94** | Male | 27 | Biopsy TLE | 0 |

After extraction, brain tissue was fixed in cold 4% paraformaldehyde (Sigma-Aldrich, St Louis, MO, USA) in 0.1M sodium phosphate buffer (PB; Panreac, 131965, Spain), pH 7.4 for 24–48h. After fixation, the tissue was washed in PB and sectioned coronally in a vibratome (150 μm thickness; Vibratome Sectioning System, VT1200S Vibratome, Leica Biosystems, Germany). Sections containing BA21 were selected and processed for Nissl-staining to determine cytoarchitecture (Fig. 1).

**Figure 1.**
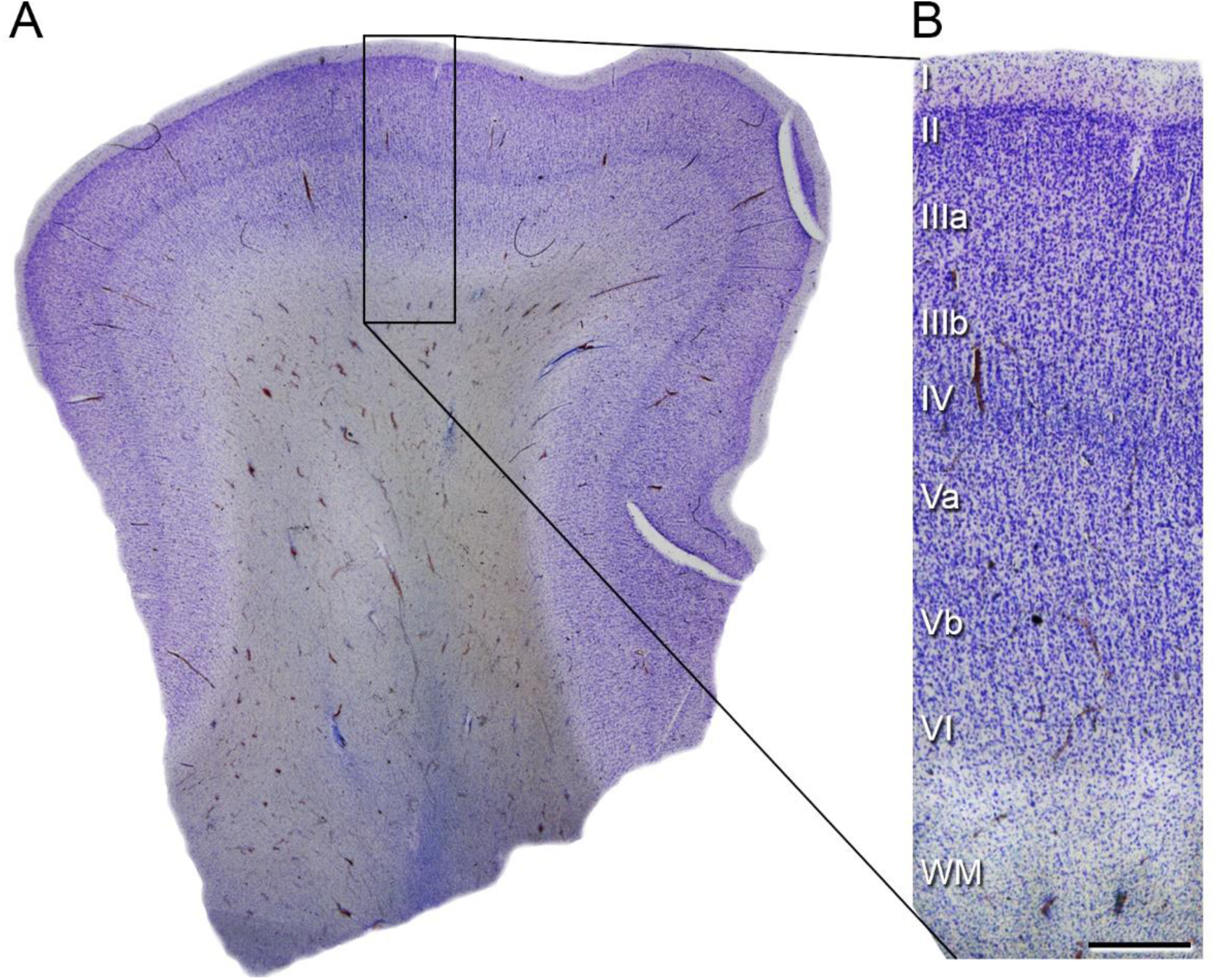
Coronal vibratome section of the human temporal neocortex. (A) Low-power photograph of a 100 µm-thick Nissl-stained section of temporal neocortex including Brodmann’s area 21 (BA21). (B) Higher magnification of the boxed area in A, showing the laminar pattern (layers I to VI are indicated). WM: white matter. Scale bar shown in B indicates 1900 µm in A and 600 µm in B.

### Electron microscopy

Sections containing BA21 were selected and postfixed for 24h in a solution containing 2% paraformaldehyde, 2.5% glutaraldehyde (TAAB, G002, UK) and 0.003% CaCl2 (Sigma, C-2661-500G, Germany) in sodium cacodylate (Sigma, C0250-500G, Germany) buffer (0.1M). The sections were treated with 1% OsO4 (Sigma, O5500, Germany), 0.1% potassium ferrocyanide (Probus, 23345, Spain) and 0.003% CaCl_2_ in sodium cacodylate buffer (0.1M) for 1h at room temperature. They were then stained with 1% uranyl acetate (EMS, 8473, USA), dehydrated and flat-embedded in Araldite (TAAB, E021, UK) for 48h at 60°C (DeFelipe and Fairén 1993). The embedded sections were then glued onto a blank Araldite block. Semithin sections (1–2 μm thick) were obtained from the surface of the block and stained with 1% toluidine blue (Merck, 115930, Germany) in 1% sodium borate (Panreac, 141644, Spain). The last semithin section (which corresponds to the section immediately adjacent to the block surface) was examined under light microscope and photographed to accurately locate the neuropil regions to be examined (Fig. 2).

**Figure 2.**
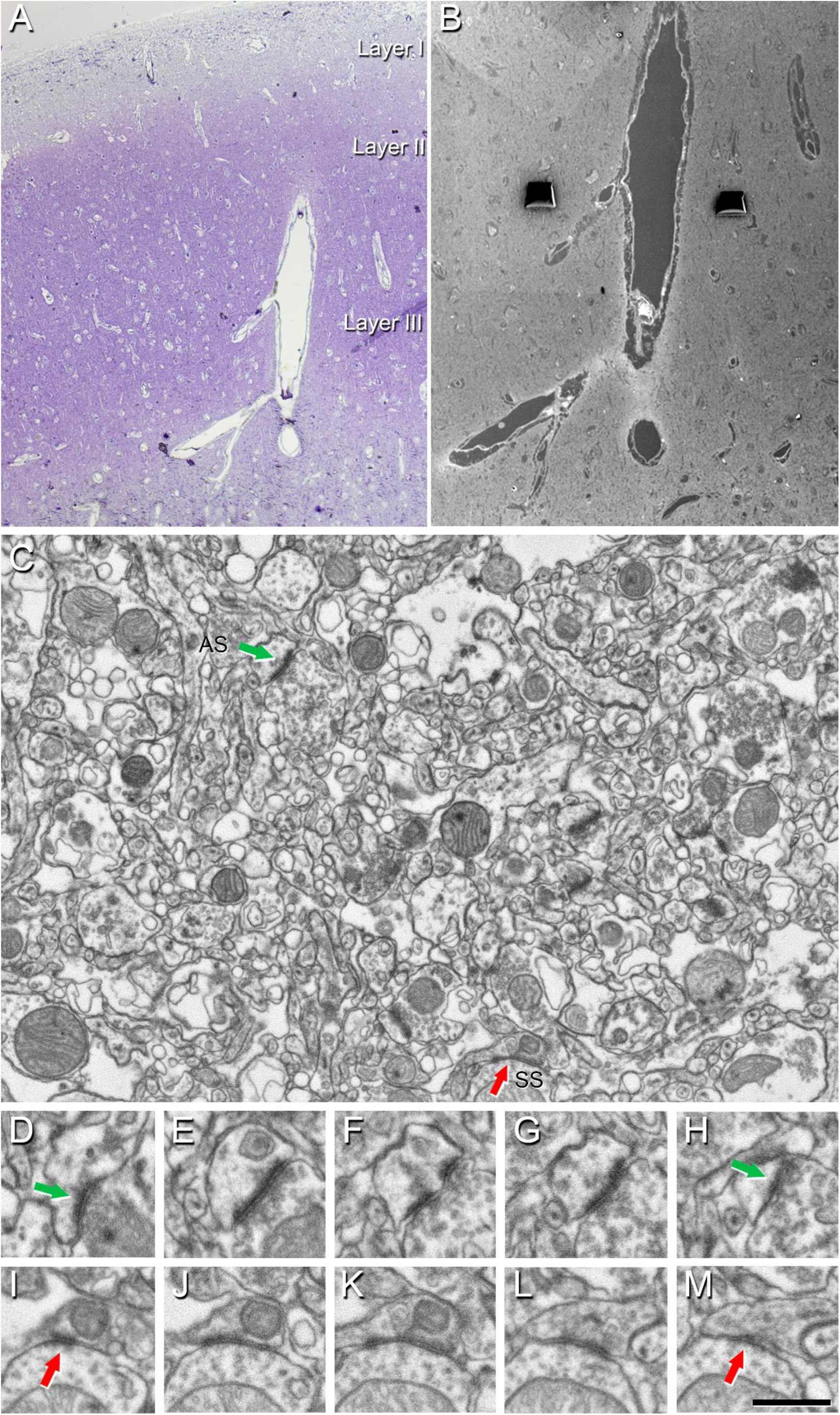
Correlative light/electron microscopy analysis of layer III of Brodmann’s area 21 (BA21). (A) Delimitation of layers is based on the staining pattern of 1 µm-thick semithin section, stained with toluidine blue, which is adjacent to the block for FIB/SEM imaging (B). (B) SEM image showing a higher magnification of the blood vessels in A to illustrate the tissue surface with the trenches made for the neuropil analyses. (C) Serial image obtained by FIB/SEM showing the neuropil of an autopsy case (AB3); two synapses are indicated as examples of asymmetric (AS, green arrow) and symmetric synapses (SS, red arrow). Synapse classification was based on the examination of the full sequence of serial images; an AS can be visualized in D–H, and an SS in I–M. See *Three-dimensional analysis of synapses* for further details. Scale bar shown in M indicates 160 µm in A, 100 µm in B, 950 nm in C, and 550 nm in D–M.

### Three-dimensional electron microscopy

The 3D study of the samples was carried out using a dual beam microscope (Crossbeam® 540 electron microscope, Carl Zeiss NTS GmbH, Oberkochen, Germany). This instrument combines a high-resolution field-emission SEM column with a focused gallium ion beam (FIB), which permits removal of thin layers of material from the sample surface on a nanometer scale. As soon as one layer of material (20 nm thick) is removed by the FIB, the exposed surface of the sample is imaged by the SEM using the backscattered electron detector. The sequential automated use of FIB milling and SEM imaging allowed us to obtain long series of photographs of a 3D sample of selected regions (Merchan-Pérez et al. 2009). Image resolution in the xy plane was 5 nm/pixel. Resolution in the z axis (section thickness) was 20 nm, and image size was 2048 x 1536 pixels. These parameters were to obtain a large enough field of view where synaptic junctions could be clearly identified, in a reasonable amount of time (approximately 12 h per stack of images).

All measurements were corrected for tissue shrinkage that occurs during processing of sections (Merchán-Pérez et al. 2009). To estimate the shrinkage in our samples, we photographed and measured the area of the vibratome sections with ImageJ (ImageJ 1.51; NIH, USA), both before and after processing for electron microscopy. The section area values after processing were divided by the values before processing to obtain the volume, area, and linear shrinkage factors (Oorschot et al. 1991), yielding correction factors of 0.90, 0.93, and 0.97, respectively. Nevertheless, in order to compare with previous studies—in which either no correction factors had been included or such factors were estimated using other methods— in the present study, we provided both sets of data. Additionally, a correction in the volume of the stack of images for the presence of fixation artifact (i.e., swollen neuronal or glial processes) was applied after quantification with Cavalieri principle (Gundersen et al. 1988b) (see Montero-Crespo et al. 2020). Every FIB/SEM stack was examined, and the volume artifact ranged from 0 to 9.4% of the volume stacks.

### Three-dimensional analysis of synapses

Stacks of images obtained by FIB/SEM were analyzed using EspINA software (EspINA Interactive Neuron Analyzer, 2.1.9; https://cajalbbp.es/espina/ Fig. 3). As previously discussed (Merchán-Pérez et al. 2009), there is a consensus for classifying cortical synapses into asymmetric synapses (AS; or type I) and symmetric synapses (SS; or type II). The main characteristic distinguishing these synapses is the prominent or thin post- synaptic density, respectively (Fig. 2). Also, these two types of synapses correlate with different functions: AS are mostly glutamatergic and excitatory, while SS are mostly GABAergic and inhibitory (DeFelipe and Fariñas 1992). Nevertheless, in single sections, the synaptic cleft and the pre- and post-synaptic densities are often blurred if the plane of the section does not pass at right angles to the synaptic junction. Since the software EspINA allows navigation through the stack of images, it was possible to unambiguously identify every synapse as AS or SS based on the thickness of the postsynaptic density (PSD) (Merchán-Pérez et al. 2009). Synapses with prominent PSDs are classified as AS, while thin PSDs are classified as SS (Gray 1959; Peters et al. 1991; Fig. 2). In addition, geometrical features —such as size and shape— and spatial distribution features (centroids) of each reconstructed synapse were also calculated by EspINA.

**Figure 3.**
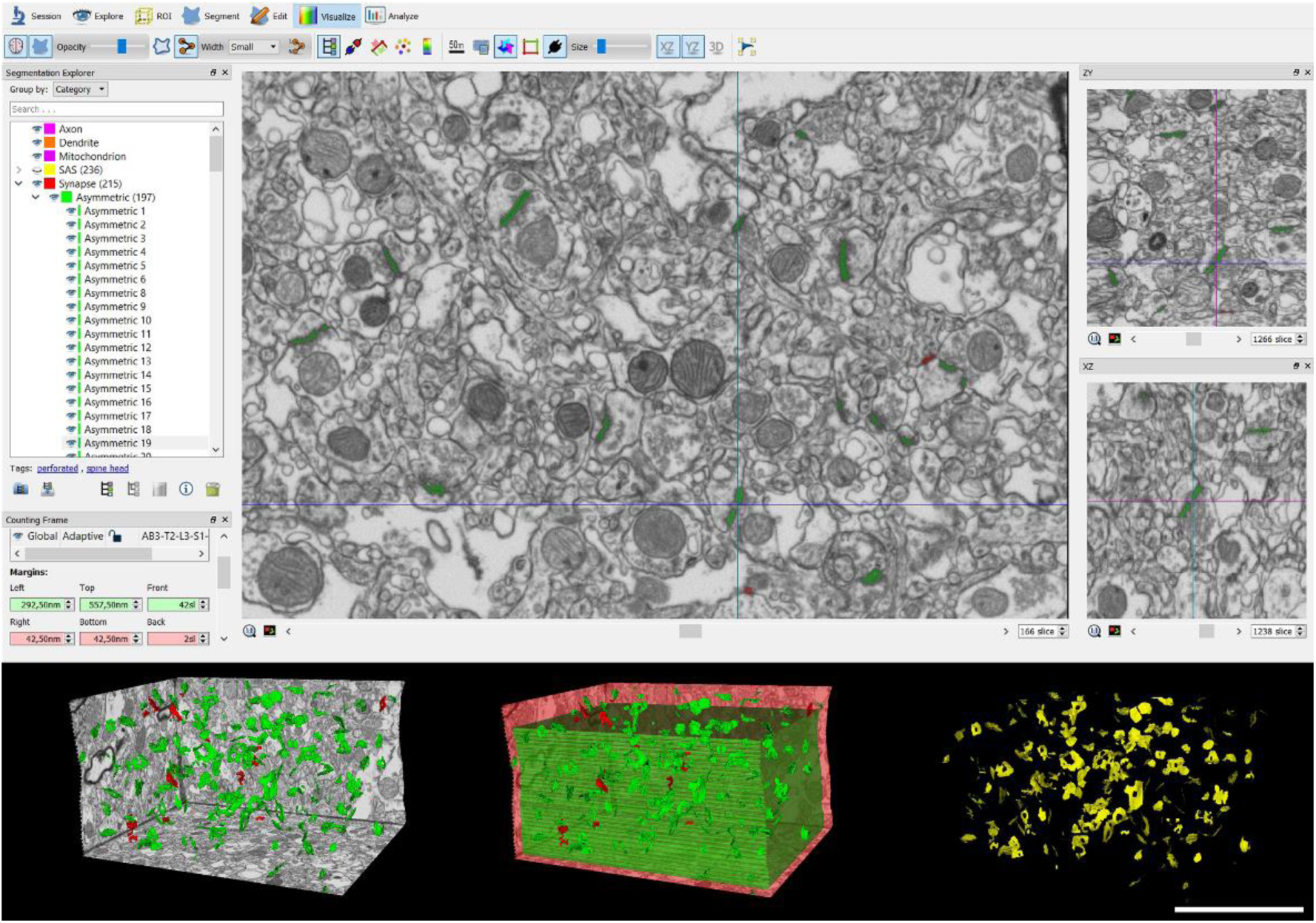
Screenshot of the EspINA software user interface. (Top) In the main window, the sections are viewed through the xy plane (as obtained by FIB/SEM microscopy). The other two orthogonal planes, yz and xz, are also shown in adjacent windows (on the right). (Bottom left) The 3D windows show the three orthogonal planes and the 3D reconstruction of AS (green) and SS (red) segmented synapses. (Bottom center) The counting frame composed by three exclusion planes (light red) and three inclusion planes (light green) with the reconstructed synapses. (Bottom right) The computed SAS for each reconstructed synapse (in yellow). Scale bar shown in the bottom right indicates 5 µm for all three of the compositions shown at the bottom.

This software also extracts the Synaptic Apposition Area (SAS) and provides its morphological measurements (Fig. 3). Given that the pre- and post-synaptic densities are located face to face, their surface areas are comparable (for details, see Morales et al. 2013). Since the SAS comprises both the active zone and the PSD, it is a functionally relevant measure of the size of a synapse (Morales et al. 2013).

EspINA was also used to visualize each of the reconstructed synapses in 3D and to detect the possible presence of perforations or deep indentations in their perimeters. Regarding the shape of the PSD, the synaptic junctions could be classified into four main categories, according to the categories proposed by Santuy et al. (2018a): macular (disk-shaped PSD); perforated (with one or more holes in the PSD); horseshoe-shaped (with an indentation) and fragmented (disk-shaped PSDs with no connection between them) (Fig. 4A).

**Figure 4.**
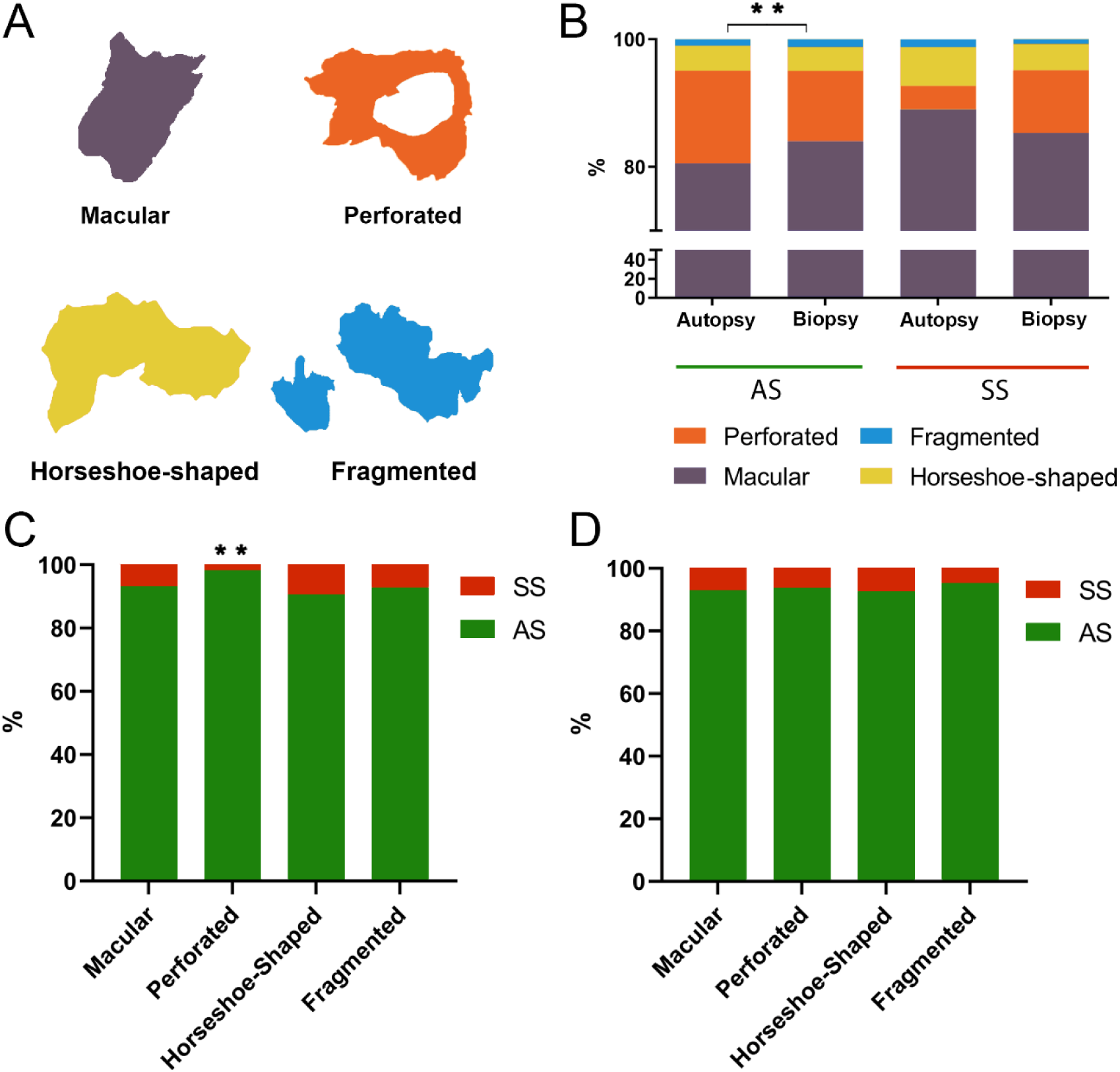
Analyses of the synaptic shape distribution in autopsy and biopsy samples from layer III of Brodmann’s area 21 (BA21). (A) Schematic representation of the shape of the synaptic junctions: macular synapses with continuous disk-shaped PSD; perforated synapses with holes in the PSD; horseshoe-shaped synapses with tortuous horseshoe-shaped perimeter with an indentation; and fragmented synapses with two or more PSDs with no connections between them. (B) Proportion of macular, perforated, horseshoe-shaped, and fragmented AS and SS in autopsy and biopsy samples. In biopsy samples, macular synapses were significantly more frequent (χ²; p=0.0054), while in autopsy samples, perforated synapses were more frequent (χ²; p=0.0013). (C) Proportion of AS and SS belonging to each morphological category in autopsy samples. The perforated synapses were significantly more frequent among AS than SS (χ²; p=0.0028). (D) Proportion of AS and SS belonging to each morphological category in biopsy samples.

To identify the postsynaptic targets of the synapses, we navigated through the image stack using EspINA to determine whether the postsynaptic element was a dendritic spine (spine, for simplicity) or a dendritic shaft. As previously described in Domínguez-Alvaro et al. (2021), unambiguous identification of spines requires the spine to be visually traced to the parent dendrite (Figs. 5 and 6; Movies 1 and 2). We refer to them as fully reconstructed spines. Additionally, when synapses were established on a spine head-shaped postsynaptic element whose neck could not be followed to the parent dendrite, we identified these elements as non-fully reconstructed spines. These non-fully reconstructed spines were identified on the basis of their size and shape, the lack of mitochondria and the presence of a spine apparatus — or because they were filled with a characteristic fluffy material (used to describe the fine and indistinct filaments present in the spines) — a term coined by Peters et al. (1991) (see also del Río and DeFelipe 1995). For simplicity, we will refer to the fully reconstructed and non-fully reconstructed spines as spines, unless otherwise specified. We also recorded the presence of single or multiple synapses on a single spine. Furthermore, we determined whether the target dendrite had spines or not.

**Figure 5.**
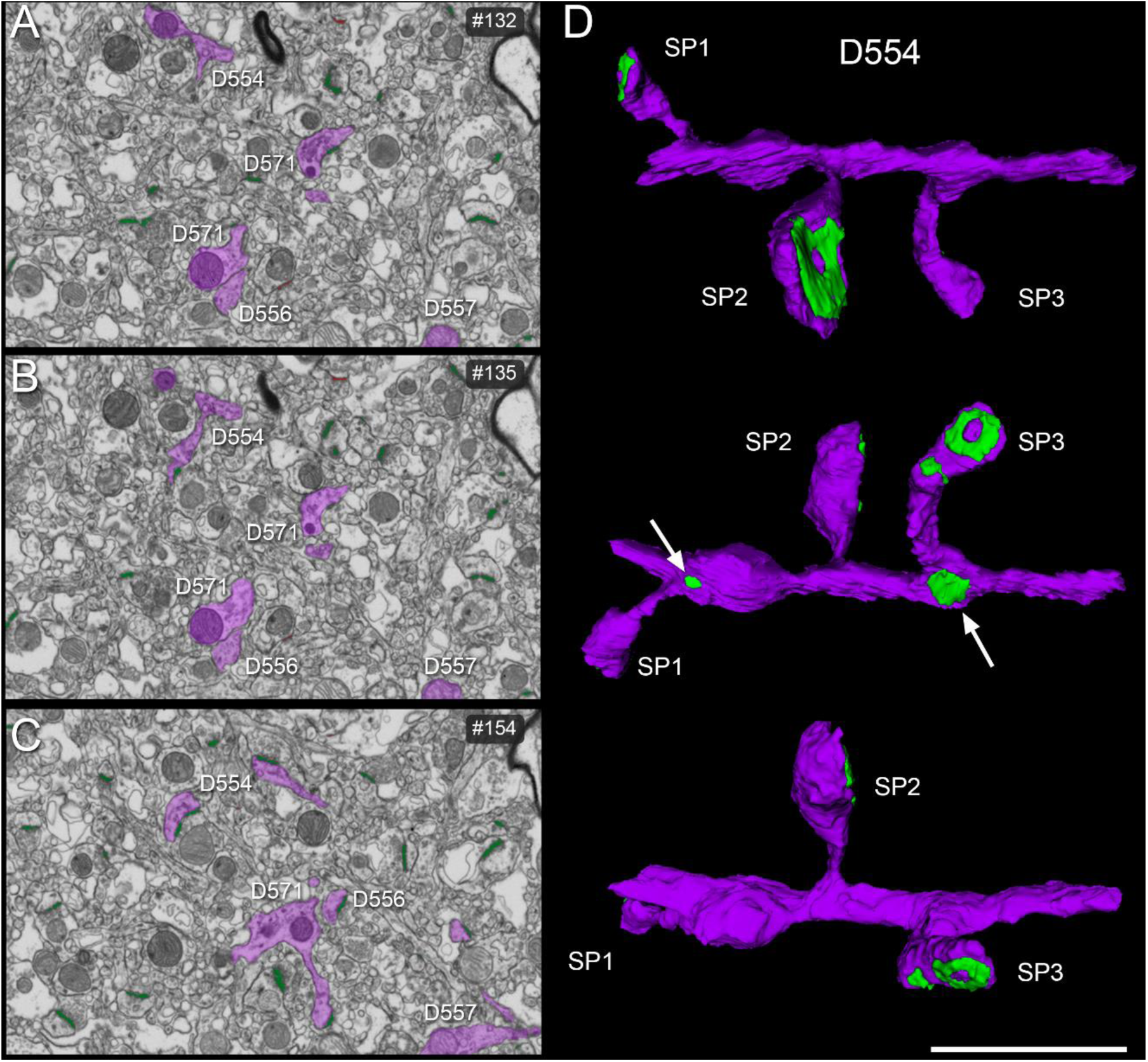
3D reconstruction of dendritic segments from FIB/SEM serial images. (A–C) Composition of electron microscopy serial images showing dendritic segments (D554, D556, D557, D571). Some reconstructions of these dendritic segments are displayed in D and Fig. 6. (D) 3D reconstructed dendritic segment D554 showing three views after rotation of the major dendritic axis at different angles. Three dendritic spines (SP) are shown, establishing asymmetric synapses (green) with different morphologies: perforated (SP2 and SP3) and macular (SP1). Arrows indicate two macular asymmetric synapses on the dendritic shaft. More examples of postsynaptic element characterization are shown in Supplementary Figures 1 and 2. Scale bar (in D) indicates 3.75 µm in A–C and 2 µm in D (See movies 1 and 2).

**Figure 6.**
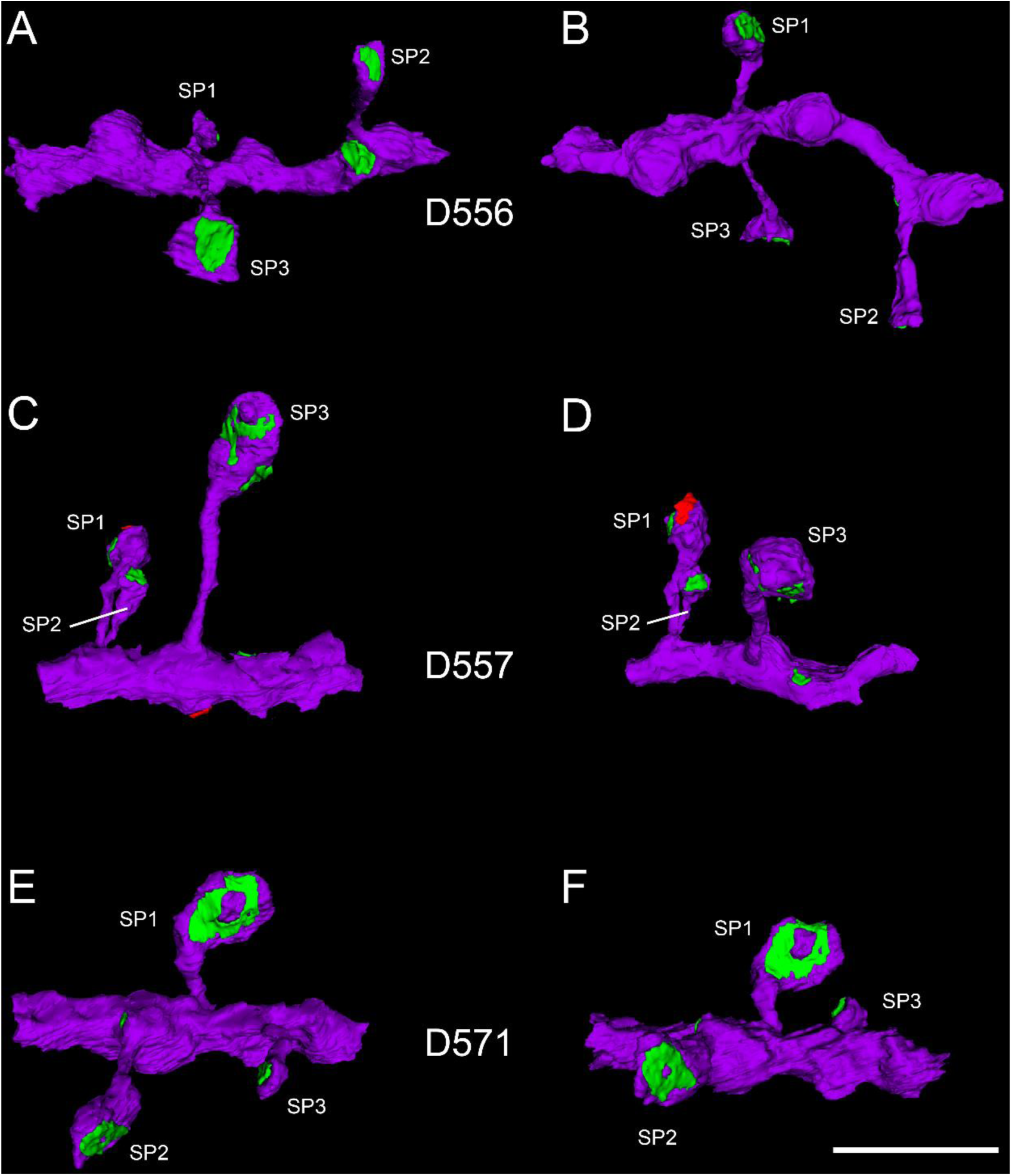
Examples of 3D reconstructed dendritic segments (purple) establishing asymmetric (green) and symmetric (red) synapses. A, B; C, D; and E, F are pairs of views after axis rotation of the dendritic segments D556 (A, B), D557 (C, D) and D571 (E, F) at different angles. Note the different sizes and shapes of the synaptic junctions. Dendritic spines SP1 and SP3 of D557 established two synapses. More examples of postsynaptic element characterization are shown in Supplementary Figures 1 and 2. Scale bar (in F) indicates 2 µm in A–F.

**Movie 1:**
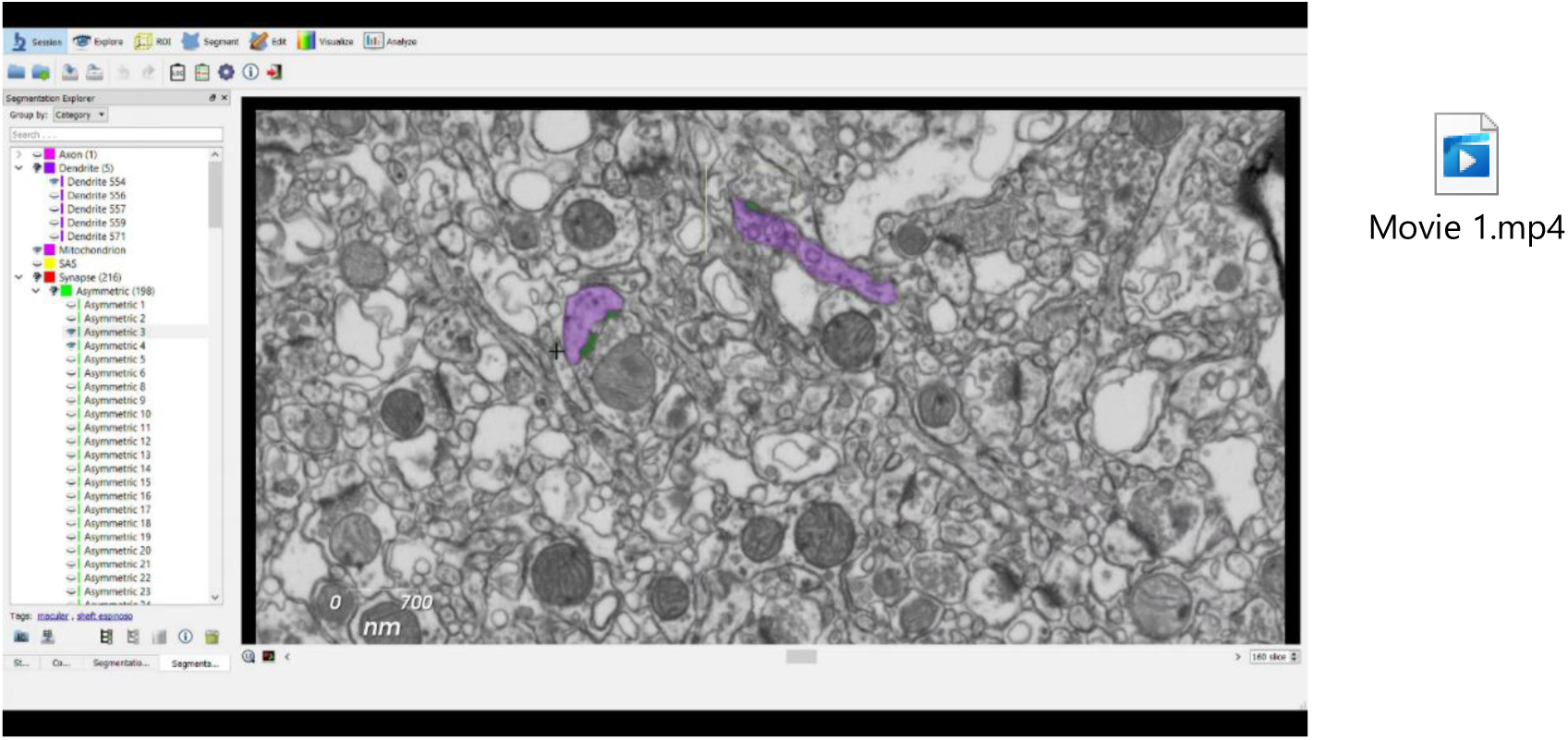
Video showing the EspINA software user interface. FIB/SEM sections are viewed through the xyz-planes. Two asymmetric (green) synapses are shown — one established on a dendritic spine and the other on the dendritic shaft (purple) of D554, also shown in Figure 5.

**Movie 2:**
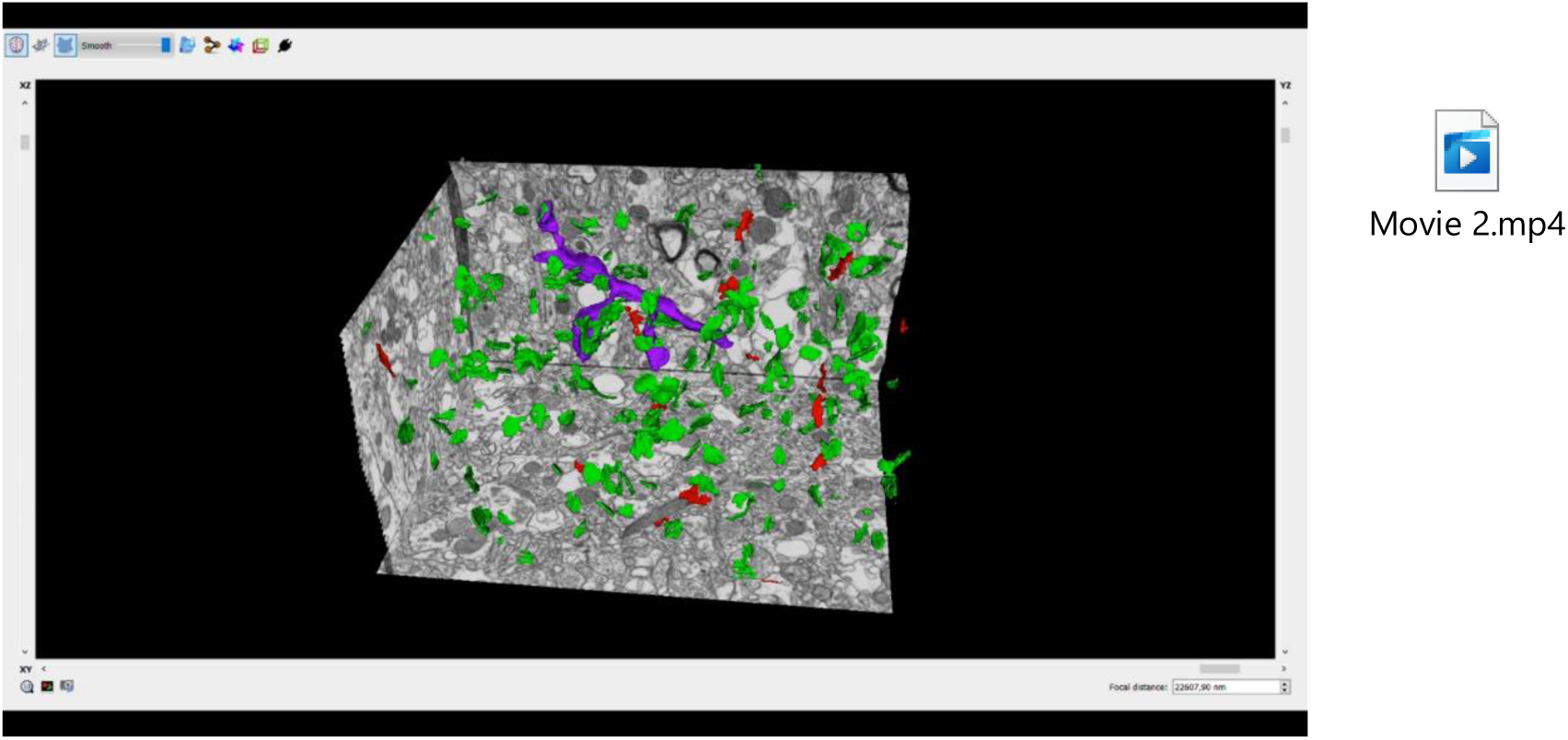
Video showing the EspINA software 3D viewer. A stack of images is fully represented in the three orthogonal planes (x, y and z). The video shows the 3D reconstruction of a single dendritic segment (purple; D554 in Figure 5) establishing synapses of different sizes and shapes and the 3D reconstruction of all asymmetric (green) and symmetric (red) synapses identified in this volume of tissue reconstructed in 3D.

### Quantification of the synaptic density

#### Direct counting method

EspINA provided the 3D reconstruction of every synapse and allowed the application of an unbiased 3D counting frame (CF). This CF is a regular rectangular prism enclosed by three acceptance planes and three exclusion planes marking its boundaries (Fig. 3). All objects within the CF are counted, as are those intersecting any of the acceptance planes, while objects that are outside the CF, or intersecting any of the exclusion planes, are not counted. Thus, the number of synapses per unit volume was calculated directly by dividing the total number of synapses counted by the volume of the CF (Merchán-Pérez et al. 2009). This method was used in all 22 stacks of images.

#### Disector method

In addition to counting directly, we also estimated the synaptic density in three stacks of images from one biopsy case (H47) using the disector method, which is based on counting the number of profiles (ΣQ−) that are present in a given section (the reference section) and that disappear in another section (the look-up section), located at a known distance in the z-axis within the unbiased CF (Gundersen 1977). The number of synapses per unit volume (NV) is calculated using the formula NV = ΣQ−/ah, where “a” is the area of the unbiased CF and “h” is the distance between the two sections. Usually after one disector is calculated, the top and bottom microphotographs are swapped and used as the new reference and look-up sections. Thus, any given pair of sections yields two estimates (Gundersen et al. 1988a). However, not all disectors are useful to estimate the number of synapses, since the distance h should not exceed 1/4 to 1/3 of the mean particle length (Gundersen et al. 1988a). Given that the mean cross-sectional length of synaptic profiles is between 280 and 350 nm (DeFelipe et al. 1999), the h value of the disectors should not exceed 70 nm if the most conservative approach is taken (1/4 of 280). In practice, we chose three times the mean section thickness (60 nm) and an unbiased CF that represent an area of 60.694 μm^2^. The variability of data obtained by the disector method tends to reduce as the number of sections sampled increases (Merchán-Pérez et al. 2009); thus, we established at least 101 pairs of disectors as a suitable value to perform the quantification.

### Spatial Distribution Analysis of Synapses

To analyze the spatial distribution of synapses, Spatial Point Pattern analysis was performed as described elsewhere (Anton-Sanchez et al. 2014; Merchán-Pérez et al. 2014). Briefly, we compared the actual position of centroids of synapses with the Complete Spatial Randomness (CSR) model — a random spatial distribution model which defines a situation where a point is equally likely to occur at any location within a given volume. For each of the 22 FIB/SEM stacks of images, we calculated three functions commonly used for spatial point pattern analysis: G, F and K functions. As described in Merchán-Pérez et al. (2014) (see also Anton-Sanchez et al. 2014) the G function, also called the nearest-neighbor distance cumulative distribution function or the event-to-event distribution, is —for a distance *d*— the probability that a typical point separates from its nearest-neighbor by a distance of *d* at the most. The F function, also known as the empty space function or the point-to-event distribution, is —for a distance *d*— the probability that the distance of each point (in a regularly spaced grid of *L* points superimposed over the sample) to its nearest synapse centroid is *d* at the most. The K function, also called the reduced second moment function or Ripley’s function, is —for a distance *d*— the expected number of points within a distance *d* of a typical point of the process divided by the intensity λ. An estimation of the K function is given by the mean number of points within a sphere of increasing radius *d* centered on each sample point, divided by an estimation of the expected number of points per unit volume. This study was carried out using the Spatstat package and R Project program (Baddeley et al. 2015).

### Statistical analysis

Statistical comparisons of synaptic density and the area of the SAS were carried out using the unpaired Mann-Whitney (MW) nonparametric U-test (normality and homoscedasticity criteria were not met). To identify possible differences regarding the area of the SAS related to its shape or its postsynaptic target, a Kruskal–Wallis (KW) nonparametric test was performed (normality and homoscedasticity criteria were not met). Frequency distribution analysis of the SAS was performed using Kolmogorov-Smirnov (KS) nonparametric test. To perform statistical comparisons of AS and SS proportions regarding their synaptic morphology and their postsynaptic target, chi-square (χ²) test was used for contingency tables. The same method was used to compare between autopsies and biopsies in relation to the synaptic type, the shape of the synaptic junctions and their postsynaptic target.

Statistical studies were performed with the GraphPad Prism statistical package (Prism 8.00 for Windows, GraphPad Software Inc., USA), Spatstat package and R Project program (Baddeley et al. 2015).

### Results

The following results were obtained in the neuropil (i.e., excluding cell bodies, blood vessels and major dendritic trunks).

### Synaptic Density

#### Direct counting method

A total of 22 stacks of images of the BA21 layer IIIA neuropil (approximately 550–750 μm from pial surface) were obtained (three stacks per case, in 6 cases; and two stacks per case, in 2 cases; total volume studied: 11 158 μm^3^) from both autopsy and biopsy samples. The number of sections per stack ranged from 259 to 317, which corresponds to a volume examined per stack ranging from 452 to 553 μm^3^ (mean: 507 μm^3^). A total of 6595 synapses were individually identified and reconstructed in 3D; of these, 4945 synapses were analyzed after discarding incomplete synapses or those touching the exclusion edges of the CF.

A total of 1337 synapses were considered for analysis from autopsy samples (total volume studied: 3659 μm^3^), and 3608 from biopsy samples (total volume studied: 7498 μm^3^) (Table 2; Supplementary table 1). The synaptic density values were obtained by dividing the total number of synapses included within the CF by its total volume. The mean synaptic density range was 0.44–0.54 synapses/μm^3^ from the autopsy samples and 0.58–0.77 synapses/μm^3^ in the case of the biopsy samples (Table 2; Supplementary table 1). The values obtained in biopsy samples were significantly higher (MW, p=0.0357) (Fig. 7).

**Figure 7.**
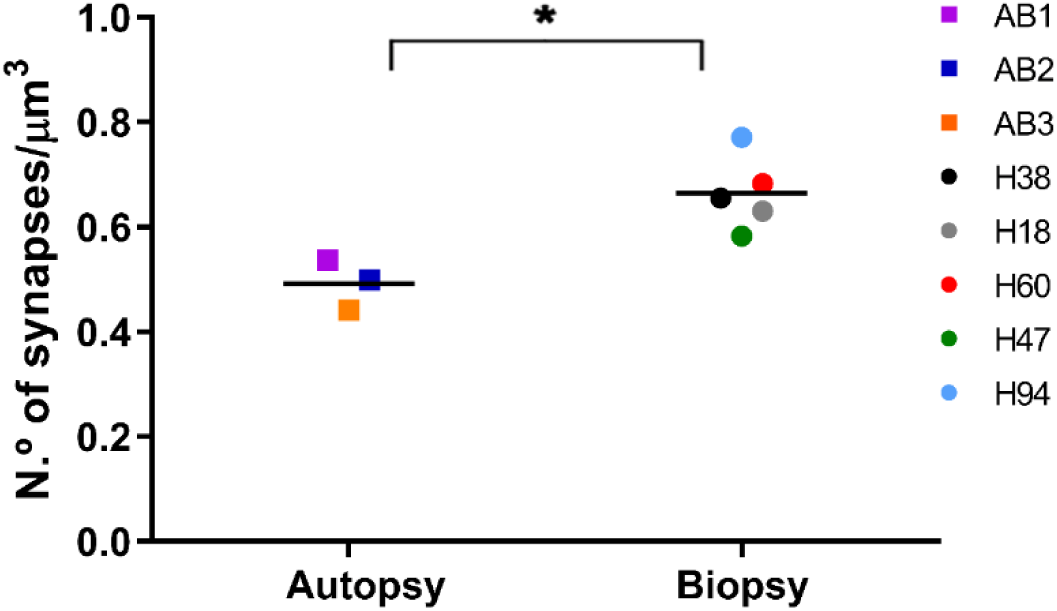
Mean synaptic density from autopsy and biopsy samples analyzed in layer III of Brodmann’s area 21 (BA21). Each symbol represents a single case according to the colored key (squares for autopsy samples, circles for biopsy samples). Significant differences in the mean synaptic density between autopsy and biopsy samples were found (MW; p=0.0357).

**Table 2.** Accumulated data obtained from the ultrastructural analysis of neuropil from layer III of BA21 in samples obtained from autopsy and biopsy. Data in parentheses have not been corrected for shrinkage. The data for individual cases are shown in Supplementary table 1. AS: asymmetric synapses; CF: counting frame; SD: standard deviation; SS: symmetric synapses.

| Group | No. AS | No. SS | No. all<br>synapses | % AS<br>(mean ± SD) | % SS<br>(mean ± SD) | CF<br>volume<br>(μm<sup>3</sup>) | No. AS<br>synapses/μm<sup>3</sup><br>(mean ± SD) | No. SS<br>synapses/μm<sup>3</sup><br>(mean ± SD) | No.<br>synapses/μm<sup>3</sup><br>(mean ± SD) | Distance to the<br>nearest neighbor<br>(nm; mean± SD) |
| --- | --- | --- | --- | --- | --- | --- | --- | --- | --- | --- |
| Autopsy | 1255 | 82 | 1337 | 93.45±2.15 | 6.55±2.15 | 2704<br>(2435) | 0.46±0.06<br>(0.51±0.06) | 0.03±0.01<br>(0.03±0.01) | 0.49±0.05<br>(0.55±0.05) | 817±51<br>(793±49) |
| Biopsy | 3363 | 245 | 3608 | 93.26±1.36 | 6.74±1.36 | 5439<br>(4898) | 0.62±0.07<br>(0.69±0.08) | 0.05±0.01<br>(0.05±0.01) | 0.66±0.07<br>(0.74±0.08) | 719±17<br>(700±17) |
| Total | 4618 | 327 | 4945 | 93.33±1.55 | 6.67±1.55 | 8143<br>(7333) | 0.56±0.10<br>(0.62±0.11) | 0.04±0.01<br>(0.04±0.01) | 0.60±0.11<br>(0.67±0.12) | 756±59 (735±56) |

#### Disector method

Quantification of the synapses using the disector method was performed to evaluate possible differences between the two different quantification methods. The disector method was used to estimate the synaptic density in the same stacks that were analyzed with the direct counting method, as well as in an additional three stacks of images from the same case (biopsy H47). A total of 319, 410 and 434 synaptic profiles were counted (ΣQ−) in FIB/SEM stacks number 1, 2 and 3, respectively. Applying the formula N_V_ = ΣQ−/ah (N_V_: number of synapses per unit volume; a: area of the unbiased counting frame; h: distance between the reference and the look-up sections), we obtained the following synaptic densities: 0.433 synapses/μm^3^, 0.557 synapses/μm^3^, and 0.590 synapses/μm^3^ in the three stacks, respectively (Supplementary Fig. 3). These values were similar to the values obtained in the same stacks by the direct counting method (Supplementary Fig. 3). That is, we did not find any difference in the synaptic densities using different estimation methods, when applied to the same samples — providing that a relatively large number of disectors were used for the estimation (101 pairs of disectors).

### Proportion of asymmetric/symmetric synapses

The proportions of AS and SS were calculated in all samples. Since synapses were fully reconstructed in 3D, it was possible to classify all synaptic contacts as AS and SS based on their postsynaptic densities (PSDs) (Fig. 2).

The proportion of AS:SS was 93:7 in both autopsy and biopsy samples (Table 2; Supplementary table 1). Hence, no significant differences in proportions between autopsy samples and biopsy samples (χ², p>0.05) were observed.

The AS:SS ratios obtained by the disector method were: 92:8, 91:9 and 89:11, respectively, in the three stacks of images analyzed. Estimation of the AS:SS ratio using the direct counting method yielded similar results — 93:7, 93:7 and 89:11, respectively.

In summary, no marked differences between the AS:SS ratios of biopsy and autopsy samples were found, regardless of which one of the two methods was used.

### Three-dimensional spatial synaptic distribution

To analyze the spatial distribution of the synapses, the actual position of each of the synapses in each stack of images was compared with a random spatial distribution model (Complete Spatial Randomness, CSR). For this, the functions G, K and F were calculated in the 22 stacks of images analyzed. One stack of images did not fit into the CSR model and it showed a slight tendency to cluster. In the remaining samples, 21 out 22 stacks, the three spatial statistical functions resembled the theoretical curve that simulates the random spatial distribution pattern, which indicated that synapses fitted a random spatial distribution model, both in the autopsy samples and the biopsy samples (Supplementary Fig. 4).

In addition, the distance of each synapse to its nearest synapse was estimated. The mean distance to its nearest neighbor measured between centroids of synaptic junctions showed significant differences (MW; p=0.0357) between autopsy and biopsy samples (817 nm and 719 nm, respectively; Table 2; Supplementary table 1), indicating that, although the synapses fitted random spatial distribution in both groups, the synapses were more separated in the autopsy samples than in the biopsy samples.

### Study of the synapse characteristics

#### Synaptic size

The study of the synaptic size was carried out analyzing the area of the SAS of each synapse identified and 3D reconstructed in all FIB/SEM stacks (Fig. 3).

In both autopsy and biopsy samples, the average size (measured by the area of the SAS) of the synapses showed significant differences (MW; p<0.0002) between AS and SS, with the latter being smaller that the former (Table 3; Supplementary table 2). These differences were also found in the frequency distribution analyses in both the autopsy samples (KS; p<0.0001) and the biopsy samples (KS; p=0.0011), again indicating that AS were larger than SS in both the autopsy (Fig. 8A) and biopsy (Fig. 8B) samples.

**Figure 8.**
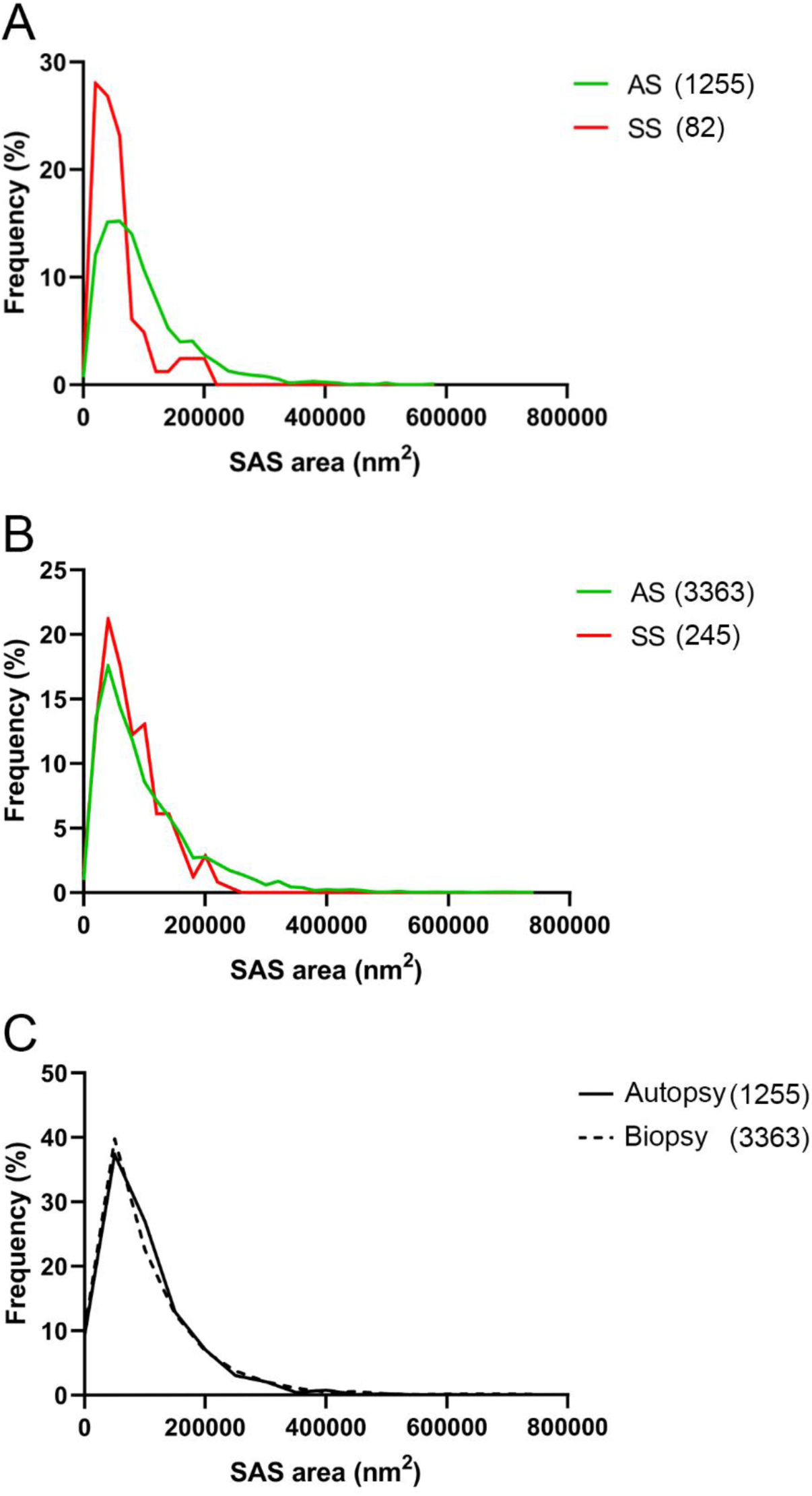
Frequency distribution of synaptic size (in terms of the SAS area). AS (green) were significantly larger than SS (red) in both autopsy samples (A; KS; p<0.0001) and biopsy samples (B; KS; p=0.0011). No significant differences were found in AS size between autopsy (continuous line) and biopsy (discontinuous line) samples (C; KS; p>0.05). Absolute numbers of analyzed synapses are shown in parentheses in the figure. AS: asymmetric synapses; SAS: synaptic apposition surface; SS: symmetric synapses.

**Table 3.** Area (nm^2^) of the SAS from autopsy and biopsy samples of layer III of BA21. All data are expressed in nm^2^ as mean±sem. Data in parentheses have not been corrected for shrinkage. The data for individual cases are shown in Supplementary table 2. AS: asymmetric synapses; SAS: synaptic apposition surface; sem: standard error of the mean; SS: symmetric synapses.

| Group | Type of synapse | Area of SAS (nm<sup>2</sup>) |
| --- | --- | --- |
| **Autopsy** | AS | 111 164±13 750 (103 383±12 787) |
|  | SS | 59 683±13 394 (55 505±12 457) |
| **Biopsy** | AS | 109 690±6356 (102 011±5911) |
|  | SS | 81 304±7878 (75 613±7327) |
| **Total** | AS | 110 243±5895 (102 526±5483) |
|  | SS | 73 196±7553 (68 072±7024) |

Upon analysis of the average area of the SAS of AS synapses in the autopsy and biopsy samples, no significant differences were found (MW; p>0.05), and the frequency distribution analyses did not reveal significant differences (KS; p>0.05; Fig. 8C). The number of SS examined in the autopsy samples was not sufficient to perform a robust statistical analysis.

#### Synaptic shape

The synapses were classified into four categories: macular (with a flat, disk-shaped PSD), perforated (with one or more holes in the PSD), horseshoe (with an indentation in the perimeter of the PSD) or fragmented (with two or more physically discontinuous PSDs) (Fig. 4A; for a detailed description, see Santuy et al. 2018a; Domínguez-Álvaro et al. 2019).

In the autopsy samples, a total of 1255 AS were identified and 3D reconstructed. The vast majority (80.6%) presented macular morphology, followed by 14.5% perforated, 3.9% horseshoe-shaped and 1% fragmented (Table 4; Supplementary table 3). For the SS, a total of 82 synapses were identified and reconstructed in 3D, and the majority also presented macular morphology (89%). Of the rest, 3.7% were perforated, 6.1% were horseshoe-shaped, and 1.2% were fragmented (Table 4; Supplementary table 3).

**Table 4.**
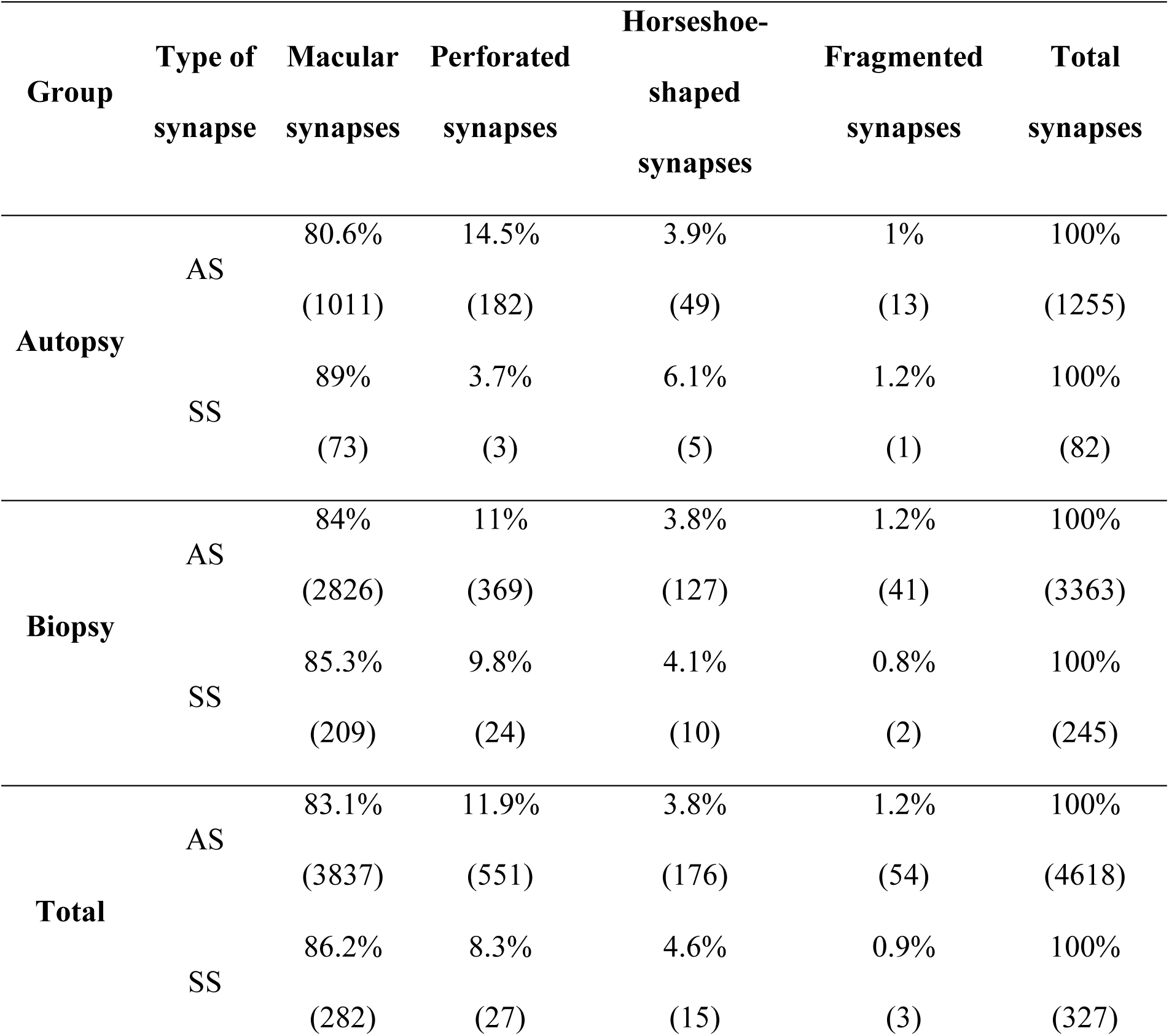
Proportion of the different shapes of synaptic junctions in the autopsy and biopsy samples from layer III of BA21. Data are given as percentages with the absolute number of synapses studied in parentheses. Data for each individual case are shown in Supplementary table 3. AS: asymmetric synapses; SS: symmetric synapses.

| Group | Type of synapse | Macular synapses | Perforated synapses | Horseshoe-shaped synapses | Fragmented synapses | Total synapses |
| --- | --- | --- | --- | --- | --- | --- |
| Autopsy | AS | 80.6% | 14.5% | 3.9% | 1% | 100% |
|  |  | (1011) | (182) | (49) | (13) | (1255) |
|  | SS | 89% | 3.7% | 6.1% | 1.2% | 100% |
|  |  | (73) | (3) | (5) | (1) | (82) |
| Biopsy | AS | 84% | 11% | 3.8% | 1.2% | 100% |
|  |  | (2826) | (369) | (127) | (41) | (3363) |
|  | SS | 85.3% | 9.8% | 4.1% | 0.8% | 100% |
|  |  | (209) | (24) | (10) | (2) | (245) |
| Total | AS | 83.1% | 11.9% | 3.8% | 1.2% | 100% |
|  |  | (3837) | (551) | (176) | (54) | (4618) |
|  | SS | 86.2% | 8.3% | 4.6% | 0.9% | 100% |
|  |  | (282) | (27) | (15) | (3) | (327) |

In the biopsy samples, a total of 3363 AS were identified and 3D reconstructed. Similarly, most of them (84%) presented macular morphology, followed by 11% perforated, 3.8% horseshoe-shaped and 1.2% fragmented (Table 4; Supplementary table 3). For the SS, a total of 245 synapses were identified and reconstructed in 3D. The majority presented macular morphology (85.3%). Of the rest, 9.8% were perforated, 4.1% were horseshoe-shaped, and 0.8% were fragmented (Table 4; Supplementary table 3).

To determine whether the shape of the synapses was related to the sample type, the frequency distributions of the morphological synaptic categories, of both AS and SS, were analyzed. Regarding AS, significant differences between biopsy and autopsy samples were found in the frequency distribution of the macular (χ², p=0.0054) and the perforated synapses (χ², p=0.0013), indicating that macular synapses were more frequent in the biopsy samples, while perforated synapses were more frequent in the autopsy samples (Fig. 4B). Concerning SS, the numbers of synapses examined separately in the autopsy and biopsy samples were not sufficient to perform a robust statistical analysis.

Regarding synaptic type (AS or SS) in the autopsy samples, from the total macular synapses, 93.3% were AS and 6.7% were SS. This proportion was maintained (χ², p>0.05) in the case of horseshoe-shaped and fragmented synapses (91:9 and 93:7 of AS:SS, respectively), but changed significantly (χ², p=0.028) in the case of the perforated synapses, which were comprised of 98.4% AS and 1.6% SS (Fig. 4C). Thus, in the autopsy samples, the ratio of AS to SS was greater in the case of perforated synapses than it was for the other morphological categories.

Regarding synaptic type (AS or SS) in the biopsy samples, from the total macular synapses, 93.1% were AS and 6.9% were SS. This proportion was maintained (χ², p>0.05) in all morphological categories (perforated: 94:6; horseshoe-shaped: 93:7; fragmented: 95:5). Thus, in the biopsy samples, no one morphological category stood out from the others as having a greater frequency of a particular synaptic type (AS or SS) (Fig. 4D).

#### Synaptic Size and Shape

We also determined whether the shape of the synapses was related to their size. For this purpose, the areas of the SAS —of both AS and SS— were analyzed according to the synaptic shape. We found that the area of the macular AS was significantly smaller than the area of the perforated, horseshoe and fragmented AS (KW, p <0.0001) in both autopsy samples (Supplementary Fig. 5A) and biopsy samples (Supplementary Fig. 5B). Concerning SS, the number of synapses examined separately in the autopsy and biopsy samples was not sufficient to perform a robust statistical analysis.

### Study of the postsynaptic elements

Postsynaptic targets were identified and classified as dendritic spines (axospinous synapses) or dendritic shafts (axodendritic synapses). We also distinguished whether the synapse was located on the neck or head of the spine (Fig. 5). When the postsynaptic element was identified as a dendritic shaft, it was classified as “with spines” or “without spines (Table 8).

**Table 8.**
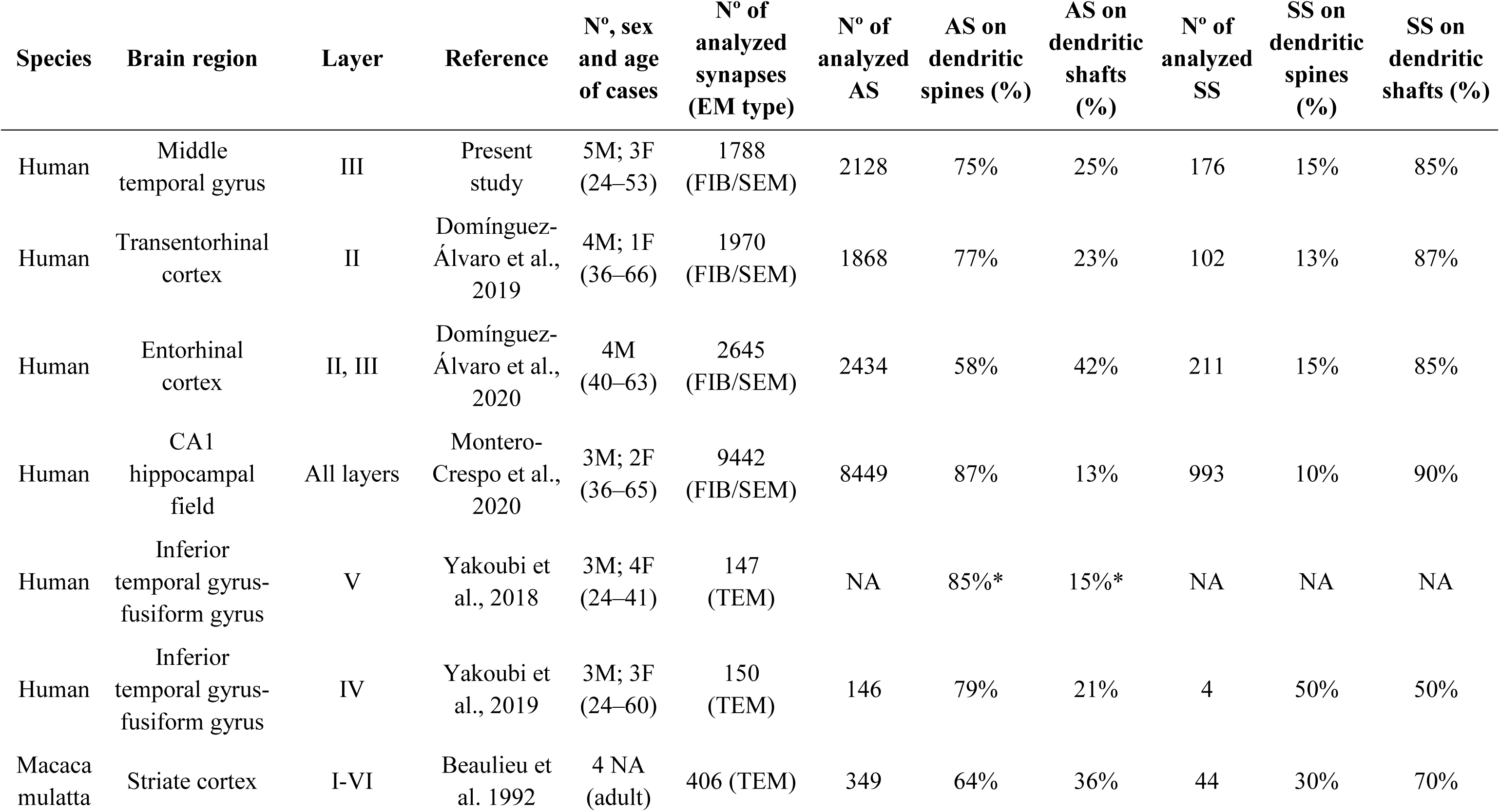
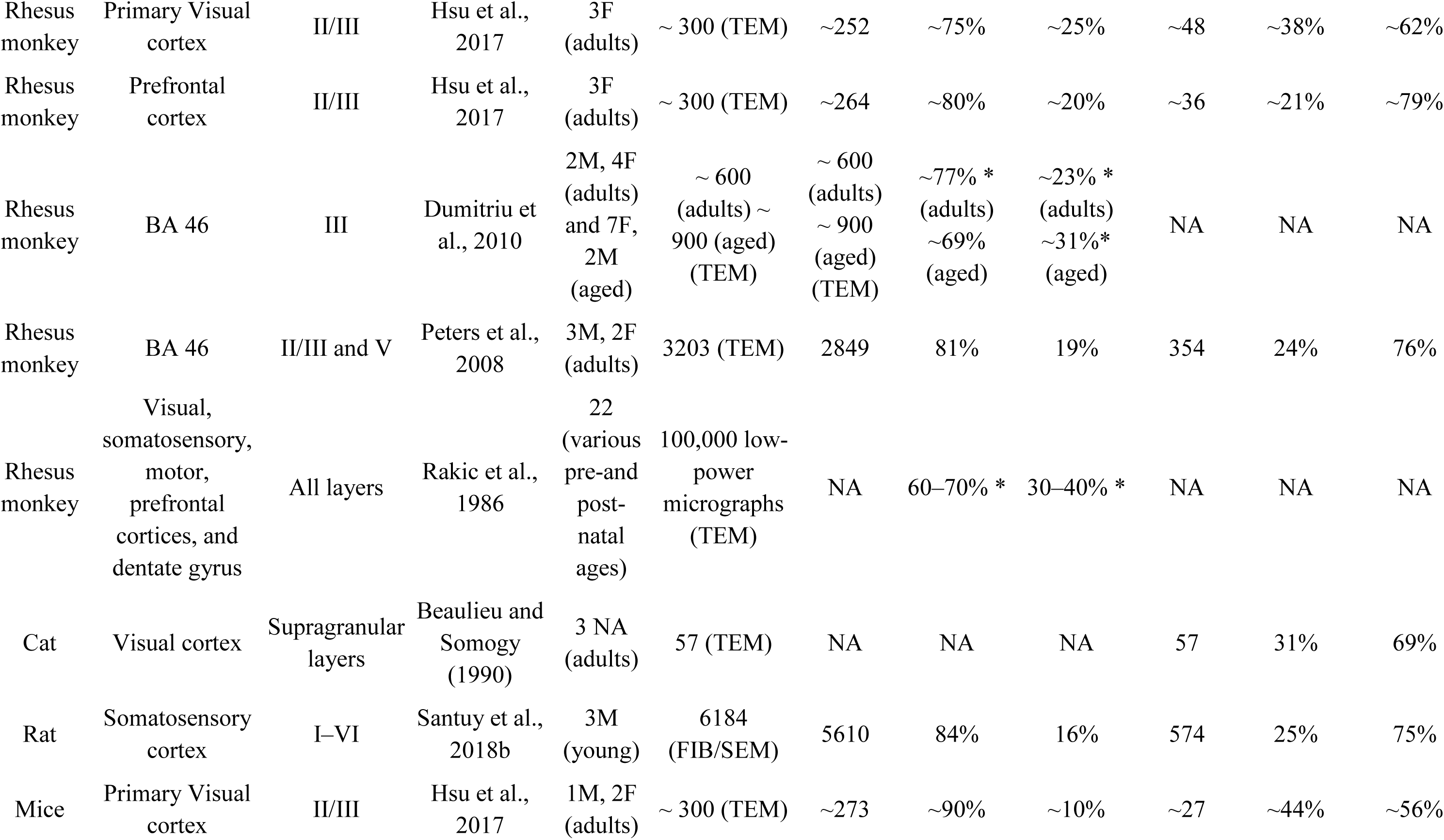
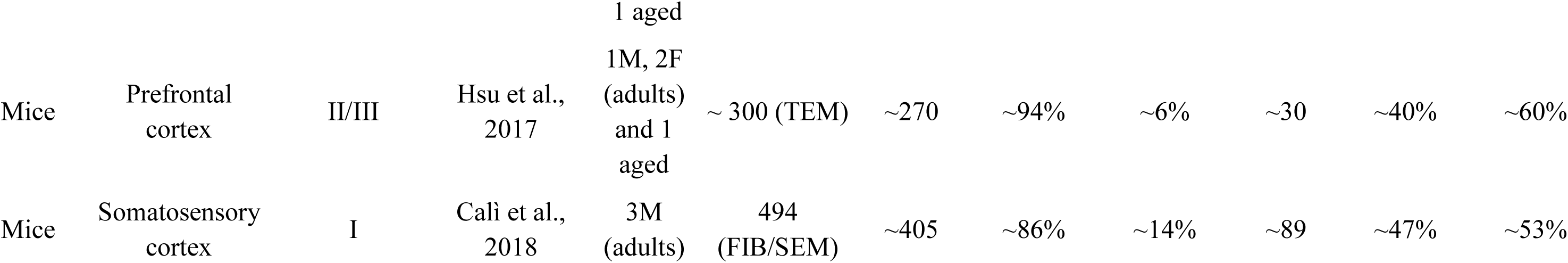
Data from the literature concerning postsynaptic target distribution in different brain species, regions, and layers. Asterisks indicate data coming from synapses in general. AS: asymmetric synapses; BA: Brodmann’s area; CA1: Cornu Ammonis 1; F: female; FIB/SEM: focused ion beam/scanning electron microscopy; M: male; NA: not available; SS: symmetric synapses; TEM: transmission electron microscopy.

In the autopsy samples, the postsynaptic elements of 1028 synapses were determined; of these, 63.62% were AS established on spine heads, 28.40% were AS on dendritic shafts, 5.45% were SS on dendritic shafts, and 1.26% were AS on spine necks (Fig. 9A). The two least frequent types of synapses were SS established on spine heads (1.07%) and SS on spine necks (0.19%) (Fig. 9A). In biopsy samples, the postsynaptic elements of 1276 synapses were determined; 71.63% of them were AS established on spine heads, 19.36% were AS on dendritic shafts, 7.29% were SS on dendritic shafts, and 0.94% were SS on spine heads (Fig. 9B). The two least frequent types of synapses were AS (0.63%) and SS (0.16%) established on spine necks (Fig. 9B).

**Figure 9.**
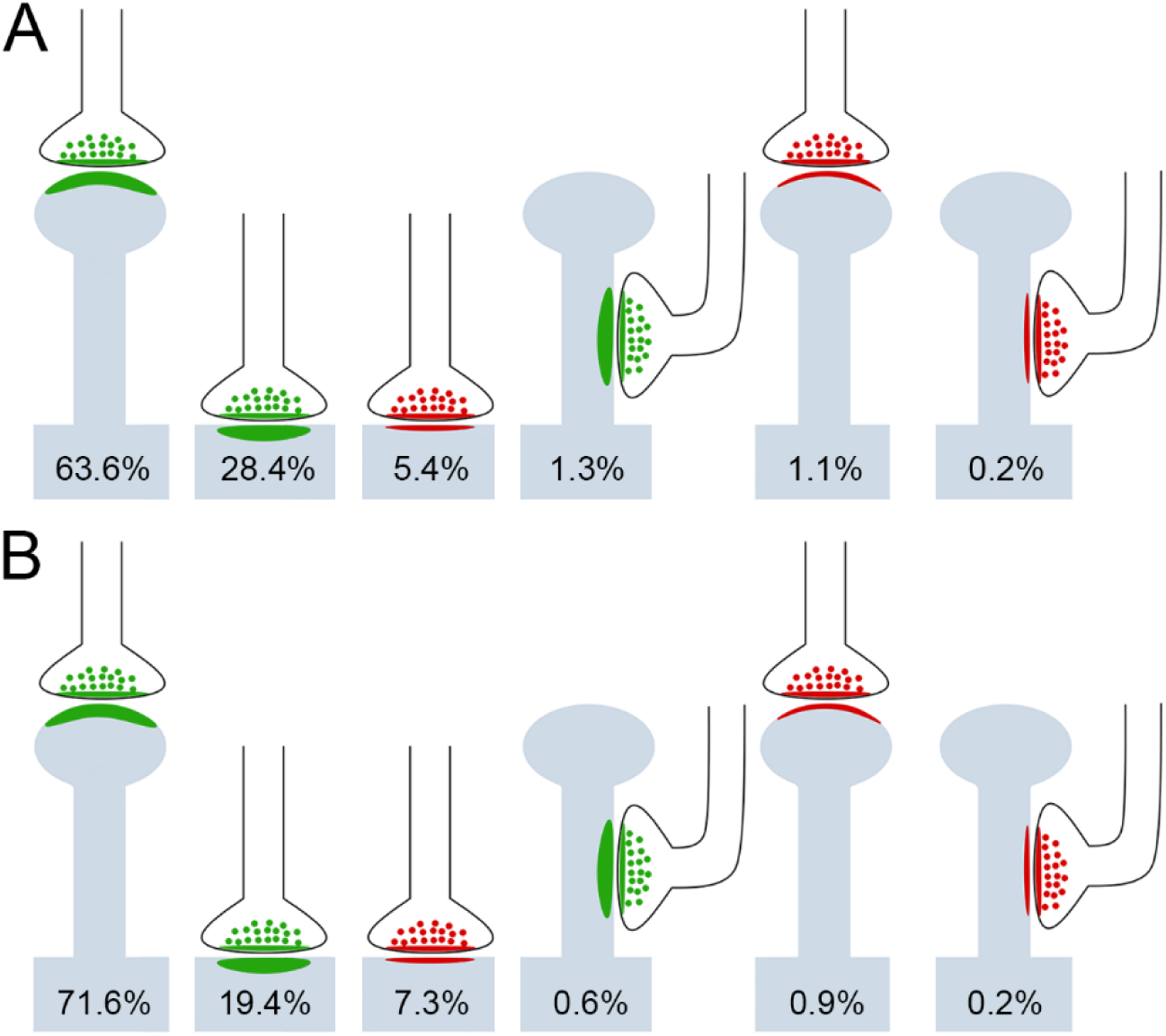
Schematic representation of the distribution of all synapses (AS and SS) on different postsynaptic targets in layer III of BA21. Spine head percentages include both fully reconstructed and non-fully reconstructed spines. Dendritic shaft percentages include dendritic shafts with and without spines. (A) Distribution of synapses on different postsynaptic targets in autopsy samples. (B) Distribution of synapses on different postsynaptic targets in biopsy samples. No significant differences were found between autopsy and biopsy samples. AS have been represented in green and SS in red. AS: asymmetric synapse; SS: symmetric synapse.

Regarding the type of synapses (AS or SS), 946 AS and 75 SS were analyzed in the autopsy samples. We found that 68.19% of the AS were established on spine heads (and 1.36% on spine necks). The remaining 30.45% of the AS were established on dendritic shafts (18.35% on dendritic shafts with spines and 12.10% on dendritic shafts without spines). In the case of the SS, 81.16% of these synapses were established on dendritic shafts (55.07% on dendritic shafts with spines and 26.09% on dendritic shafts without spines), while 15.94% were established on spine heads — and the remaining 2.90% were found to be on spine necks (Table 5; Supplementary table 4; Fig. 10).

**Figure 10.**
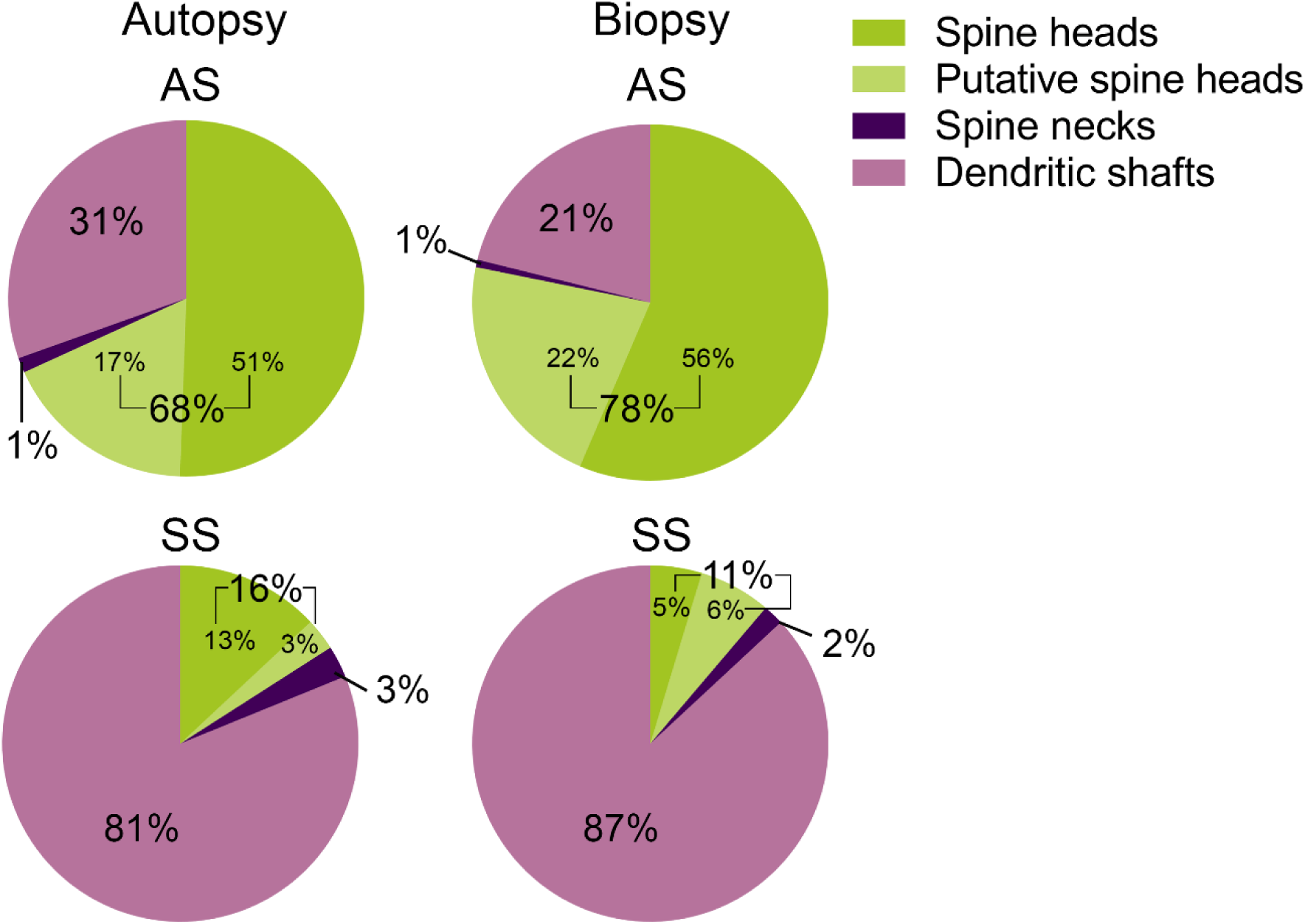
Proportions of postsynaptic targets of AS and SS in layer III of BA21. In both autopsy and biopsy samples, asymmetric synapses and symmetric synapses show significant preference for spine head targets (including both fully reconstructed and non- fully reconstructed spines) and dendritic shaft targets, respectively (χ²; p<0.0001). Regarding AS, biopsy samples showed a greater proportion of spine head targets than autopsy samples (χ²; p<0.0001). AS: asymmetric synapse; SS: symmetric synapse.

**Table 5.**
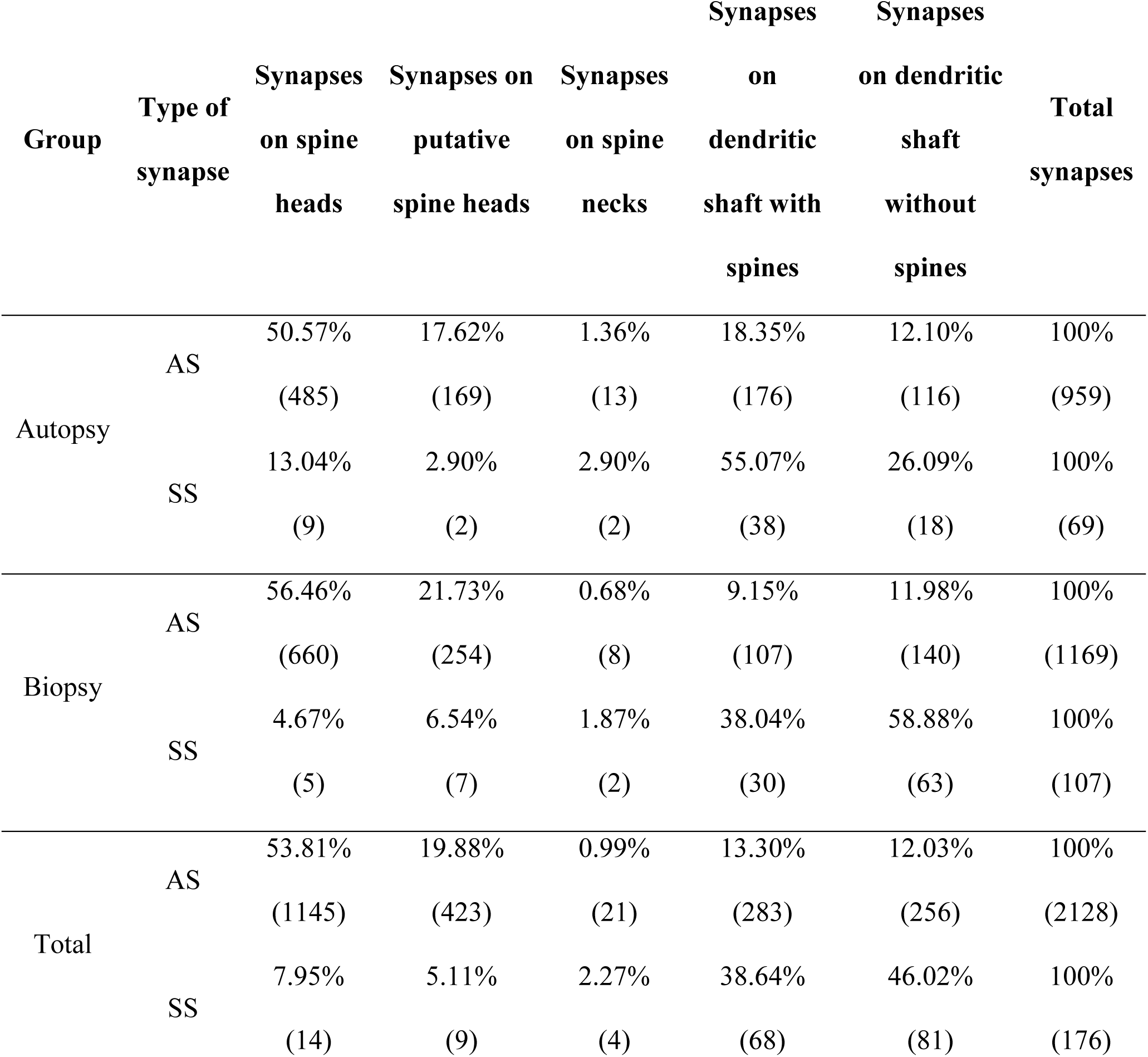
Distribution of AS and SS on spines and dendritic shafts in autopsy and biopsy samples from layer III of BA21. Synapses on spines have been sub-divided into those established on spine heads that are fully reconstructed or non-fully reconstructed and those established on spine necks. Moreover, we differentiated between dendritic shafts with spines and without spines. Data are given as percentages with the absolute number of synapses studied in parentheses. Data for each individual case are shown in Supplementary table 4. AS: asymmetric synapses; SS: symmetric synapses.

In the biopsy samples, the postsynaptic elements of 1169 AS and 107 SS were determined. We found that 78.19% of the AS were established on spine heads (considering spine heads and putative spine heads together) and 0.68% were on spine necks. The remaining 21.13% of the AS were established on dendritic shafts (9.15% on dendritic shafts with spines and 11.98% on dendritic shafts without spines). Regarding the SS, 86.92% were established on dendritic shafts (28.04% on dendritic shafts with spines and 58.88% on dendritic shafts without spines), while 11.21% were established on spine heads — and the remaining 1.87% were found to be on spine necks (Table 5; Supplementary table 4; Fig. 10).

Considering all types of synapses established on the spine heads, the proportion of AS:SS was 98:2 and 99:1, in autopsy and biopsy samples, respectively; while in those established on dendritic shafts, this proportion was 84:16 (autopsy) and 73:27 (biopsy). Since the overall AS:SS ratio was 93:7, the present results show that AS and SS did show a ‘preference’ for a particular postsynaptic element; AS showed a preference for the heads of the spines (χ², p<0.0001), while the SS showed a preference for the dendritic shafts (χ², p<0.0001), (Table 5; Fig. 10).

To determine whether there was a difference between the different postsynaptic elements found in the autopsy and the biopsy samples, frequency distribution of the postsynaptic elements, of both AS and SS, were analyzed. Significant differences (χ², p<0.0001) between the autopsy and biopsy samples were found with regard to the AS spine heads and the dendritic shafts frequencies — spine heads were more frequent in the biopsy samples, while dendritic shafts were more frequent in the autopsy samples (Table 5; Fig. 10). Concerning SS, the number of synapses examined was not sufficient to perform a robust statistical analysis.

To detect the presence of multiple synapses, an analysis of the spine heads was performed to determine the number and type of synapses established on them. The most frequent finding was a spine head establishing a single AS (97.0% and 97.8% in the autopsy and biopsy samples, respectively), followed by two AS (1.4% and 0.9% in the autopsy and biopsy samples, respectively), while the proportion of spine heads receiving one AS and one SS was 0.9% (autopsy samples) and 0.5% (biopsy samples). The presence of a single SS on a spine head was very low indeed (0.8% in both the autopsy and biopsy samples; Fig. 11). The analysis of the synapses established on spine heads to detect possible differences in the presence of multiple synapses between autopsies and biopsies was discarded due to the low numbers of SS established on spine heads.

**Figure 11.**
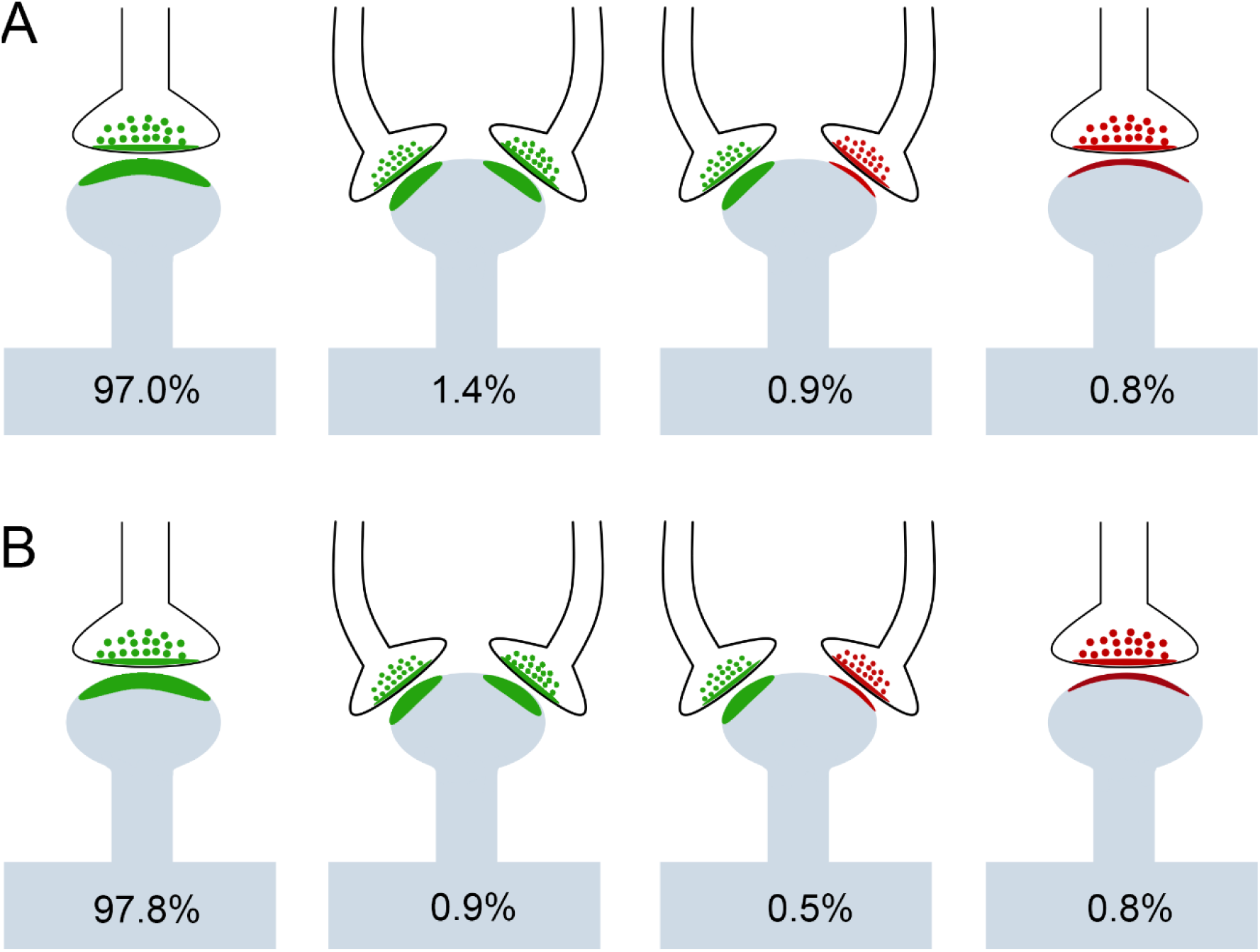
Schematic representation of single and multiple synapses on dendritic spine heads (including both fully reconstructed and non-fully reconstructed spines) in autopsy (A) and biopsy (B) samples. Percentages of each type are indicated. AS have been represented in green and SS in red. AS: asymmetric synapse; SS: symmetric synapse.

#### Postsynaptic elements and synaptic size

Finally, we analyzed whether there was a relationship between the synaptic size and the type of postsynaptic element. This study was carried out using the data of the area of the SAS of each synapse whose postsynaptic element had been identified as a spine head, spine neck, or a dendritic shaft with or without spines. Synapses established on non-fully reconstructed spines were discarded. AS established on spine heads were larger than AS established on spine necks in both the autopsy (KW; p=0.0023) and biopsy samples (KW; p=0.0244). Also, AS established on spine heads were larger than AS established on dendritic shafts in both autopsy (KW, p=0.0002) and biopsy samples (KW; p<0.0001) (Supplementary Fig. 6). Concerning SS, the number of synapses was not sufficient to perform a robust statistical analysis.

## Discussion

We have performed a detailed description of the synaptic organization of the neuropil of layer III from BA21 using FIB/SEM in human tissue from both autopsy and biopsy samples. Critical data on synaptic structure and functionality can be obtained from determination of the synaptic density, AS:SS ratio, and 3D spatial distribution, as well as the shape and size of the synapses and their postsynaptic targets. The present results provide a new, large, quantitative ultrastructure dataset in 3D of the synaptic organization of this particular brain region.

The following major results were obtained: (i) the estimated synaptic density values were lower in autopsy samples than in biopsy samples (ii) the AS:SS ratios were similar in the autopsy and biopsy samples; (iii) in all samples, the synapses fitted into a random spatial distribution; (iv) the area of the AS was larger than that of SS, in both the autopsy and the biopsy samples; (v) most synapses displayed a macular shape — smaller than complex-shaped synapses; (vi) most AS were established on dendritic spines, while most SS were established on dendritic shafts; (vii) AS on spine heads were more frequent in the biopsy samples, while dendritic shafts were more frequent in the autopsy samples.

### Synaptic density

Synaptic density is a critical parameter when it comes to describing the synaptic organization of a particular brain region, in terms of connectivity and functionality. The mean synaptic density found in layer III of the human BA21 was 0.61 synapses/μm^3^, but differences were found between the autopsy and biopsy samples, with lower values observed in autopsy samples (0.49 synapses/μm^3^) than in biopsy samples (0.66 synapses/μm^3^). Previous EM studies on the human temporal neocortex have shown either higher synaptic density values in layers II/III (0.73 synapses/μm^3^; Davies et al. 1987) and in layer III (0.99 synapses/μm^3^; Alonso-Nanclares et al. 2008)— or lower values in other layers (0.35 and 0.32 synapses/μm^3^; Huttenlocher and Dabholkar 1997; Tang et al. 2001; Table 6). A recent study performed in layer IV of the human temporal neocortex reported a synaptic density of 0.002 synapses/μm^3^ (Yakoubi et al. 2019), which is an extremely low value considering the present results and those reported in the literature (Table 6).

**Table 6.**
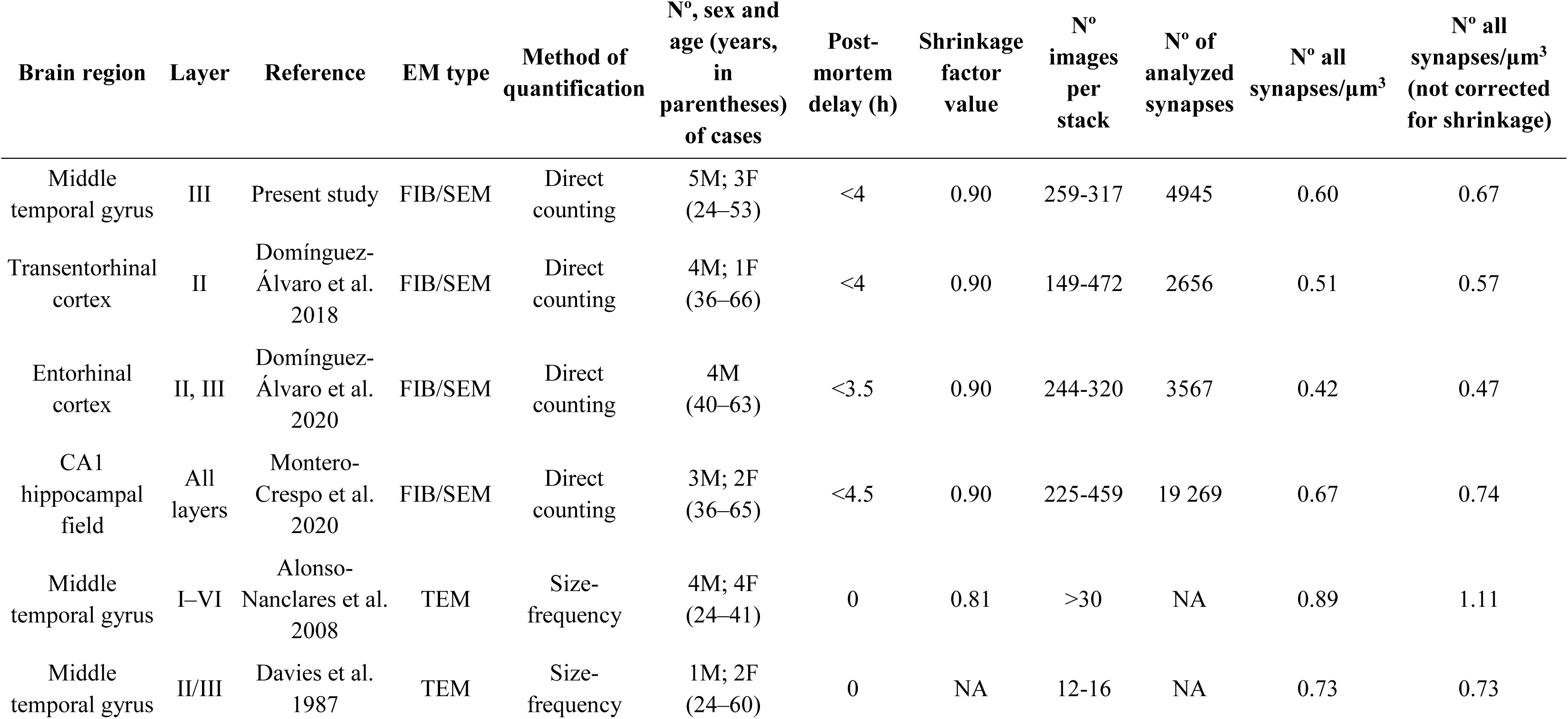
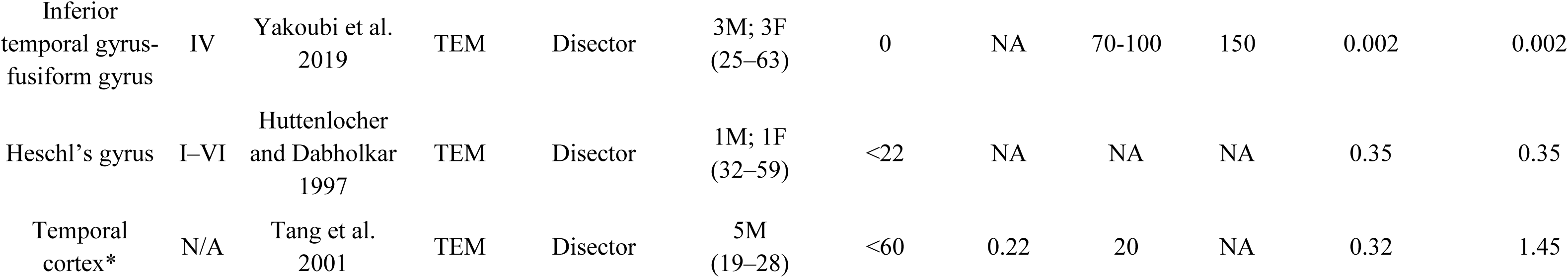
Data obtained from the present study and from the literature concerning the synaptic density in the human temporal cortex, including the hippocampal formation. Note that the analysis carried out by FIB/SEM technology provides data coming from larger volumes of examined tissue than TEM. Asterisk indicates that no data regarding the temporal region examined was provided. BA: Brodmann’s area; CA1: Cornu Ammonis 1; F: female; FIB/SEM: focused ion beam/scanning electron microscopy; M: male; NA: not available; TEM: transmission electron microscopy.

| Brain region | Layer | Reference | EM type | Method of quantification | Nº, sex and age (years, in parentheses) of cases | Post-mortem delay (h) | Shrinkage factor value | Nº images per stack | Nº of analyzed synapses | Nº all synapses/$\mu\text{m}^3$ | Nº all synapses/$\mu\text{m}^3$ (not corrected for shrinkage) |
| --- | --- | --- | --- | --- | --- | --- | --- | --- | --- | --- | --- |
| Middle temporal gyrus | III | Present study | FIB/SEM | Direct counting | 5M; 3F (24–53) | <4 | 0.90 | 259-317 | 4945 | 0.60 | 0.67 |
| Transentorhinal cortex | II | Domínguez-Álvaro et al. 2018 | FIB/SEM | Direct counting | 4M; 1F (36–66) | <4 | 0.90 | 149-472 | 2656 | 0.51 | 0.57 |
| Entorhinal cortex | II, III | Domínguez-Álvaro et al. 2020 | FIB/SEM | Direct counting | 4M (40–63) | <3.5 | 0.90 | 244-320 | 3567 | 0.42 | 0.47 |
| CA1 hippocampal field | All layers | Montero-Crespo et al. 2020 | FIB/SEM | Direct counting | 3M; 2F (36–65) | <4.5 | 0.90 | 225-459 | 19 269 | 0.67 | 0.74 |
| Middle temporal gyrus | I–VI | Alonso-Nanclares et al. 2008 | TEM | Size-frequency | 4M; 4F (24–41) | 0 | 0.81 | >30 | NA | 0.89 | 1.11 |
| Middle temporal gyrus | II/III | Davies et al. 1987 | TEM | Size-frequency | 1M; 2F (24–60) | 0 | NA | 12-16 | NA | 0.73 | 0.73 |
| Inferior temporal gyrus-fusiform gyrus | IV | Yakoubi et al. 2019 | TEM | Disector | 3M; 3F (25–63) | 0 | NA | 70-100 | 150 | 0.002 | 0.002 |
| Heschl's gyrus | I–VI | Huttenlocher and Dabholkar 1997 | TEM | Disector | 1M; 1F (32–59) | <22 | NA | NA | NA | 0.35 | 0.35 |
| Temporal cortex* | N/A | Tang et al. 2001 | TEM | Disector | 5M (19–28) | <60 | 0.22 | 20 | NA | 0.32 | 1.45 |

Differences in the estimated synaptic densities (Table 6) may stem from multiple factors, including technical issues affecting the preservation of the ultrastructure. Fixation techniques affect the preservation of the ultrastructure, and therefore the estimated synaptic density may be different from one study to the next. In the present study, although we used the same fixative solution, biopsies were fixed immediately after removal, while tissue from the autopsies was fixed up to 4h post-mortem. A long- postmortem delay for the fixation of brain tissue samples can interfere with synaptic identification (Glausier et al. 2019). Additionally, the dimensions of the brain tissue blocks in biopsies are generally smaller than those from autopsies, which could also influence the penetration of the fixative, such that the fixation in smaller blocks is better (Glausier et al. 2019). The above-mentioned reports from Huttenlocher and Dabholkar (1997) and Tang et al. (2001) used brain tissue samples with a postmortem delay of up to 22 hours and 60 hours, respectively. Therefore, the final values of the synaptic density might be lower due to a lack of proper preservation and identification of the synapses. Moreover, some studies (e.g., Davies et al. 1987; Huttenlocher and Dabholkar 1997) did not take into account that estimations, including morphological measurements, are affected by the tissue shrinkage caused by tissue preparation for electron microscopy, which may modify the final estimates. In the present study, we have applied a correction factor to all lineal, surface and volume measurements to overcome the shrinkage derived from electron microscopy processing in chemical-fixed tissue. However, we can not address how the particular subcellular elements are affected by the chemical fixation. Electron microscopy studies based on cryo-fixed samples seem to be the best option to represent the living state of the tissue. Using this cryo-fixation technique, larger volumes of extracellular space compared to the chemical-fixed samples are reported, and some subcellular elements such as the dendritic spine necks (but not the heads) are shrunken to a greater degree when compared to chemical fixation (Korogod et al. 2015, Tamada et al. 2020). However, as discussed in Tamada et al. (2020), the volumes of well cryo-preserved tissue are still far lower than the volumes that can currently be preserved with chemical fixation, so there are still many limitations with regard to performing a large ultrastructural and quantitative analysis using cryo-fixation techniques. Nevertheless, caution is required when extrapolating the geometrical measurements of ultrastructural elements obtained by chemical fixation to the living state. In the present study, we have applied a correction factor to all lineal, surface and volume measurements to overcome the shrinkage derived from electron microscopy processing in chemical-fixed tissue. However, we can not address how the particular subcellular elements are affected by the chemical fixation. Electron microscopy studies based on cryo-fixed samples seems to be the best option to represent the living state of the tissue. Using this cryo fixation technique, an appreciable amounts of extracellular space, that not appears in the chemical-fixed samples, is showed, and some subcellular elements such the dendritic spines necks are differently shrunken when comparing to chemical fixation (Korogod et al. 2015, Tamada et al. 2020). However, the volumes of well cryo-preserved tissue is still far from the currents volumes that we can preserve with chemical fixation, so there are still many restrictions to perform a large ultrastructural and quantitative analyses using this fixation technique (Tamada et al. 2020). Nevertheless, caution must be necessary to extrapolate the geometrical measurements of ultrastructural elements obtained by chemical fixation to the living state.

Moreover, a lower synapse number has been associated with normal aging in several human neocortical regions (Adams 1987; Benavides-Piccione et al. 2013; Mohan et al. 2018), including the temporal cortex (Berchtold et al. 2013; Henstridge et al. 2016). In our study, the autopsy samples came from individuals aged 45, 53 and 53 years old, whereas the individuals that provided the biopsy samples were younger (24, 27, 31 36, and 41 years old). Therefore, the lower synaptic density observed in autopsy samples (0.49±0.05 mm^3^) compared to biopsy samples (0.66±0.07 mm^3^) could be related to a combined effect of the fixation techniques and the possible influence of normal aging. Perhaps more importantly, the differences in the estimated synaptic density reported in the present study compared with that observed in the previous reports may stem from the quantification method and the volume of tissue examined. In some of the above-mentioned TEM studies, synaptic density was calculated using estimations based on the number of synaptic profiles per unit area (“size-frequency method”) in single ultrathin sections of tissue, whereas —in others— the “disector method” in several consecutive ultrathin sections was applied. That is, the number of synapses per volume unit was estimated from two-dimensional samples. However, both the size-frequency and the disector methods are usually applied on a relatively small number of sections due to the difficulties associated with obtaining large numbers of serial sections. As has been shown previously by Merchán-Pérez et al. (2009) and confirmed in the present study, these stereological methods are accurate but only when applied to a large number of sections. Yakoubi et al. (2019) used the disector method on a large number of excellent-quality TEM serial sections (70–100 sections) from human biopsies belonging to three women and three men, with ages ranging from 25 to 63 years old, with fixation times of 0 hours (Table 6). Thus, their samples might be comparable to ours in terms of ultrastructure preservation and age (Table 1). However, the synaptic density reported by Yakoubi et al. (2019) was more than one hundred times lower than our present results and any previously reported values in the human cerebral cortex, including the hippocampal formation (Davies et al. 1987; Huttenlocher and Dabholkar 1997; Tang et al. 2001; Alonso-Nanclares et al. 2008; Domínguez-Álvaro et al. 2020; Montero-Crespo et al. 2020).

To further examine the influence of the choice of estimation method, we compared the values obtained by direct counting and by the disector method in three stacks of FIB/SEM images. With both methods, we obtained comparable values when all the serial images were used for the estimations (0.64 synapses/μm^3^ by direct counting, and 0.53 synapses/μm^3^ by the disector method; Supplementary Fig. 3). Indeed, the larger the number of serial sections analyzed, the lower the variability in the final synaptic density estimations. In addition, analysis of 30 TEM images of the same case and layer using the size-frequency method revealed a much higher density (1.04 synapses/μm^3^; Alonso-Nanclares et al. 2008). Summing up, a notable difference in the synaptic density values can be observed depending on the method applied and the number of serial sections used. Previous 3D ultrastructural studies using FIB/SEM in the same autopsy cases as those analyzed in the present study have shown that the synaptic density in other cortical regions —such as layer II of the transentorhinal cortex (0.51 synapses/μm^3^; Domínguez-Álvaro et al. 2018) and layers II and III of the entorhinal cortex (0.42 synapses/μm^3^; (Domínguez-Álvaro et al. 2020)— is within the range of the present results. However, estimations in the human hippocampal CA1 field in these autopsy cases have shown that synaptic density ranged from 0.45 to 0.99 synapses/μm^3^ (in the stratum oriens and in the superficial part of the stratum pyramidale, respectively) (Montero-Crespo et al. 2020). Since the processing and analysis methods were identical, similarities (to the entorhinal cortices) and differences (with respect to the CA1 superficial pyramidal layer) may be attributable to the specific characteristics of the layer and brain region analyzed, highlighting the fact that synaptic density greatly depends on the human brain layer and/or region being studied (reviewed in DeFelipe 2015).

### Proportion of synapses and spatial synaptic distribution

It has been established, in different brain regions and species, that the cortical neuropil has a larger proportion of asymmetric synapses than symmetric synapses (reviewed in DeFelipe et al. 2002; DeFelipe 2011). Previous studies have shown that the percentage of AS and SS varies —80–95% and 20–5%, respectively— in all the cortical layers, cortical areas and species examined so far by TEM (Beaulieu and Colonnier 1985; Megías et al. 2001; Bourne and Harris 2011; DeFelipe 2011, 2015).

In the present study, the AS:SS ratio was 93:7 in both autopsy and biopsy samples, and no differences were found between the two kinds of brain tissue. Similar results have been found using the same methods in the human entorhinal cortex (92:8; Domínguez-Álvaro et al. 2020), the human transentorhinal cortex (96:4; Domínguez-Álvaro et al. 2018) and in the human CA1 hippocampal field (95:5, except in the stratum lacunosum moleculare, in which the AS:SS ratio was 89:11; Montero-Crespo et al. 2020). Thus, it would appear that the present AS:SS ratio data provide among the highest and lowest proportions previously observed in different cortical regions using TEM, for AS and SS, respectively.

The significance of this relatively high proportion of AS and low proportion of SS, and the invariance of this proportion in the human cerebral cortex, is difficult to interpret due to the differences in the cytoarchitecture, connectivity and functional characteristics of different brain regions.

Since the present results showed similar AS:SS ratios in the neuropil of the above-mentioned brain regions, it could be interpreted that the AS:SS ratio in the dendritic arbor of the different neurons may be similar. However, in other cortical regions of a variety of species, it has been shown that there are differences in the number of inputs of GABAergic and glutamatergic synapses in several neuronal types (e.g., DeFelipe and Fariñas 1992; Freund and Buzsáki 1996; DeFelipe 1997; Somogyi et al. 1998; Schubert et al. 2007; Markram et al. 2015; Tremblay et al. 2016). Thus, it would be necessary to examine the synaptic inputs of each particular cell type to determine possible differences in their AS:SS ratio, even though the overall AS:SS proportion does not vary in the neuropil.

Regarding the spatial organization of synapses, we found that the synapses were randomly distributed in the neuropil samplesfrom both autopsies and biopsies. This type of spatial distribution has also been found in a number of regions of the human brain (frontal cortex, transentorhinal cortex, entorhinal cortex and CA1 hippocampal field; (Blazquez-Llorca et al. 2013; Domínguez-Álvaro et al. 2018, 2020; Montero-Crespo et al. 2020). Therefore, the present results regarding the random spatial distribution of synapses and the AS:SS proportion (ranging from 92:8 to 96:4) support the idea of widespread ‘rules’ of the synaptic organization of the human cerebral cortex.

### Synaptic size and shape

It has been proposed that synaptic size correlates with release probability, synaptic strength, efficacy and plasticity (Nusser et al. 1998; Kharazia and Weinberg 1999; Takumi et al. 1999; Ganeshina et al. 2004a; Tarusawa et al. 2009; Matz et al. 2010; Holderith et al. 2012; Südhof 2012; Montes et al. 2015; Biederer et al. 2017). Several methods have traditionally been used to estimate the size of synaptic junctions (Table 7).

**Table 7.**
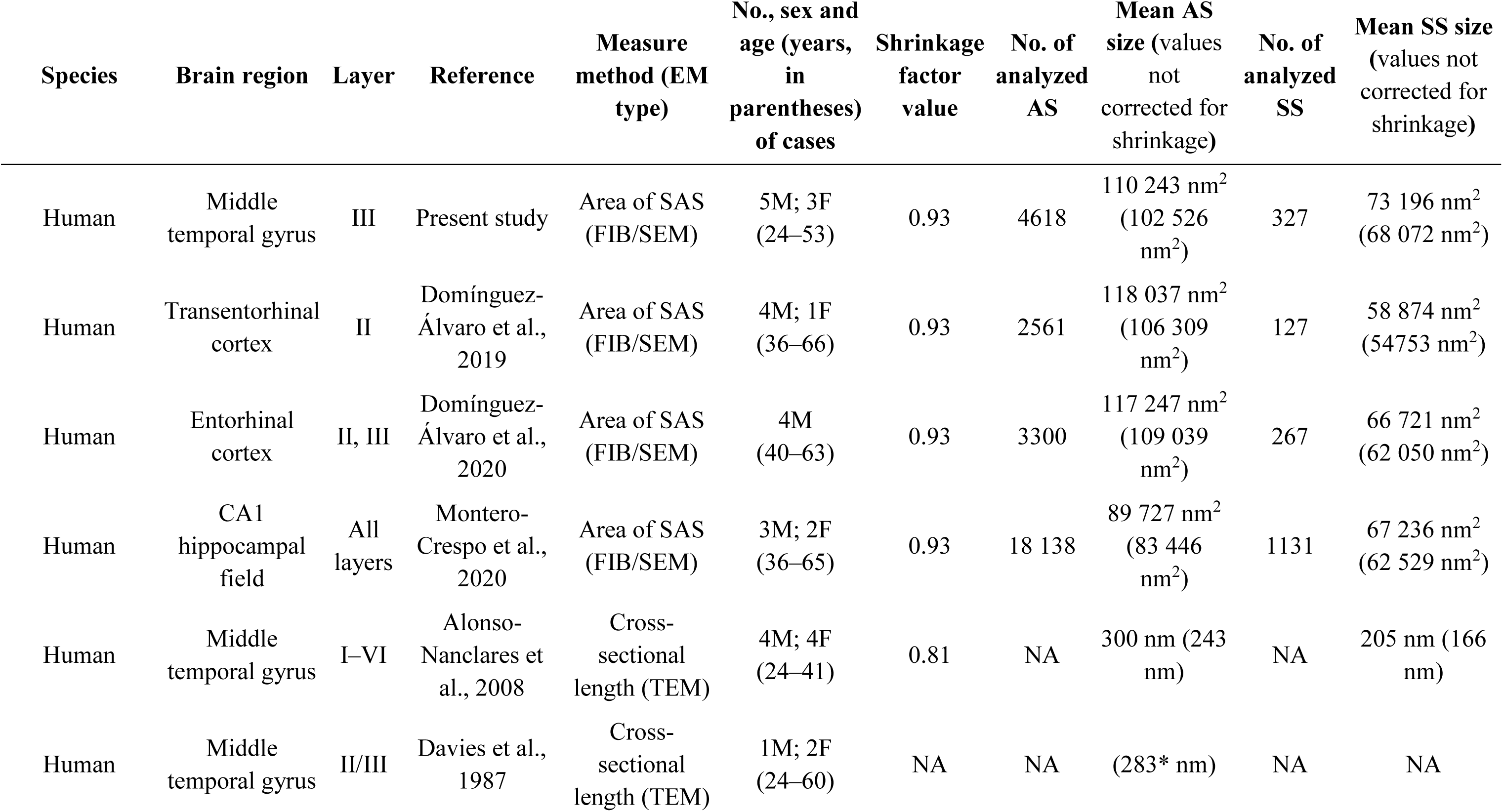
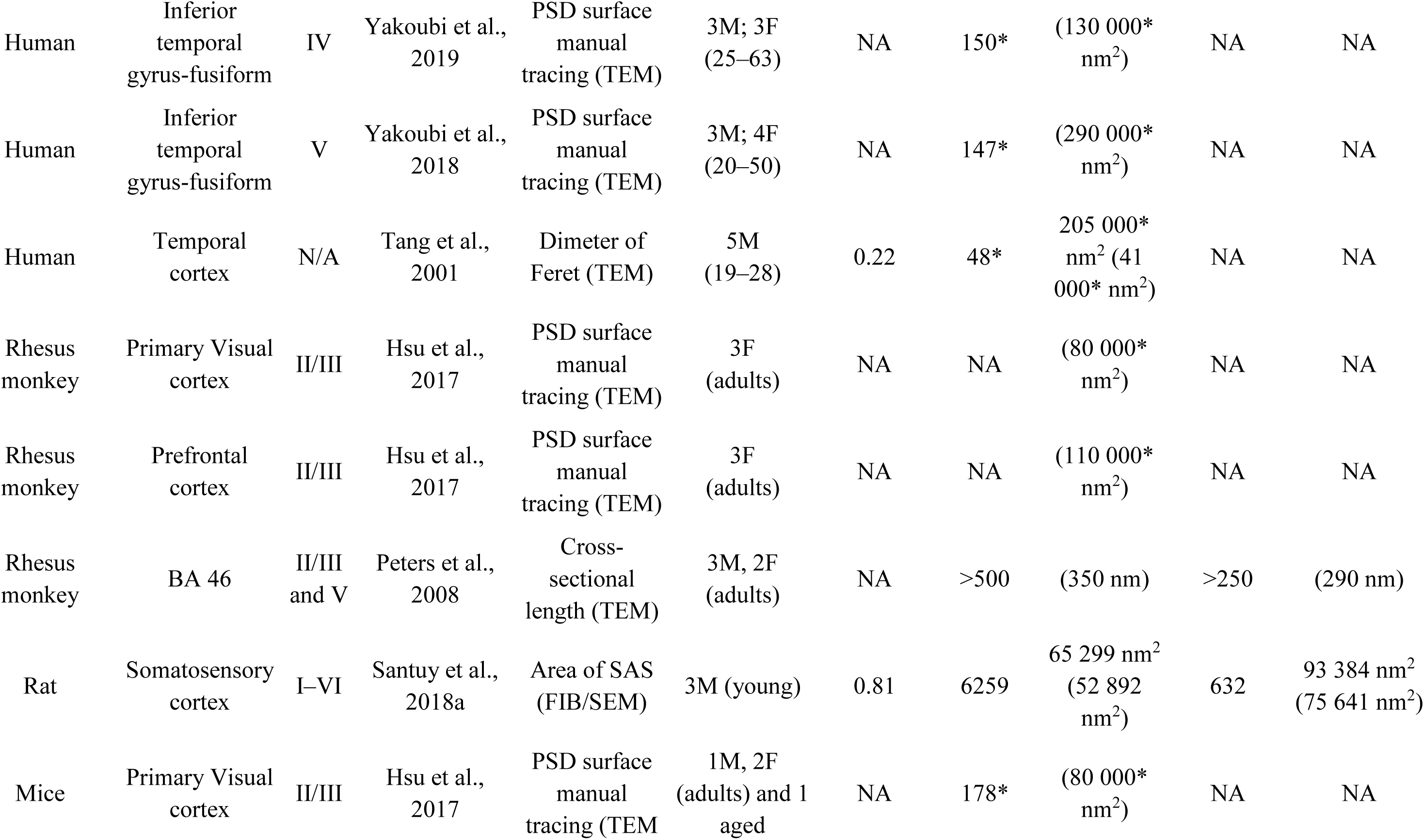
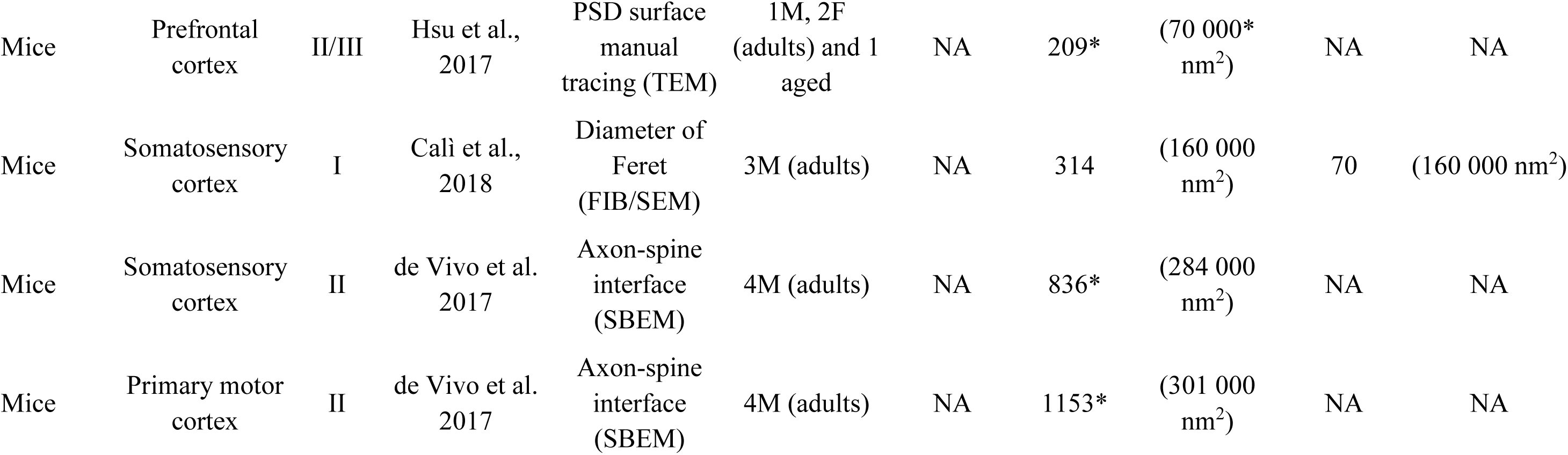
Data from the present results and from literature concerning synaptic size in different brain species, regions, and layers. Asterisks (*) indicate data coming from synapses in general (that is, without specifying whether they are AS or SS). BA: Brodmann’s area; CA1: Cornu Ammonis 1; F: female; FIB/SEM: focused ion beam/scanning electron microscopy; M: male; NA: not available; SBEM: serial block-face scanning electron microscopy; TEM: transmission electron microscopy.

The simplest of these methods is to measure the cross-sectional length of synaptic junctions in TEM micrographs (Davies et al. 1987; Alonso-Nanclares et al. 2008). This method has obvious limitations since it is based on individual 2D images where a portion of synapses cannot be fully characterized (DeFelipe et al. 1999), and it also reduces size estimation to a 1D measurement. Alternatively, the PSD can be reconstructed from the series of sections and its contour can be measured in 3D (Hsu et al. 2017; Yakoubi et al. 2018, 2019). Another measurement that has been used to estimate the size of synaptic junctions in 3D is the Feret diameter, which is equivalent to the diameter of the smallest sphere circumscribing the reconstructed object (Merchán-Pérez et al. 2014; Calì et al. 2018). However, this does not accurately describe the morphology of synapses, since it obviously oversimplifies the geometric characteristics of the object measured. Another indirect measurement of the size of the synaptic junction is the axon-spine interface, which represents the total apposition surface between the membrane of the axonal bouton and the membrane of the dendritic spine (de Vivo et al. 2017). Although the area of the axon-spine interface and the PSD are correlated (Cheetham et al. 2014), data coming from the axon-spine interface are not comparable with ours, because the SAS is always inside the axon-spine interface and thus it is smaller than the axon-spine interface. Therefore, we have used the SAS, which is equivalent to the interface between the AZ and the PSD. Moreover, our methodology provides information on the shape of the PSD, as well as information about synapses established on dendritic shafts, which cannot be obtained from axon-spine interface measurements.

There are very few studies on human brain that provide data regarding the size of the synaptic junctions. For example, using manual PSD tracing reconstruction in layer IV and V of the temporal neocortex, Yakoubi et al. (2018, 2019) reported that the mean size of the PSD was 130 000 nm^2^ and 290 000 nm^2^ in layer IV and V, respectively. The values obtained in layer IV were similar to our measurements for the area of the SAS of AS in layer IIIA (110 243 nm^2^), and also similar to our previous results in layer II of the human transentorhinal cortex (118 037 nm^2^; Domínguez-Álvaro et al. 2019) and layers II and III of the entorhinal cortex (110 311 nm^2^ and 124 183 nm^2^, respectively; Domínguez-Álvaro et al. 2020). Thus, it would be necessary to analyze SAS in all cortical layers of BA21 to determine whether layer V has the largest PSD and whether other layers have SAS that are similar to —or smaller than— those found in layers IIIA and IV.

Regarding other species, the synaptic size found in the present study was similar to those described by manual PSD reconstruction in the prefrontal cortex and larger than in the primary visual cortex of the rhesus monkey (110 000 and 80 000 nm^2^, respectively (Hsu et al. 2017). The synaptic sizes observed in our results were also larger than those described in rat somatosensory cortex with the same method (65 299 nm^2^; Santuy et al. 2018a). Different synaptic sizes have also been reported in mice with different methods; using manual PSD reconstruction, Hsu et al. (2017) reported a mean synaptic size of 80 000 nm^2^ and 70 000 nm^2^ in the primary visual cortex and prefrontal cortex, respectively. Using the Feret diameter, Calì et al. (2018) found a mean synaptic size of 160 000 nm^2^, while using axon-spine interface measurements, de Vivo et al. (2017) reported values of 284 000 nm^2^ in the somatosensory cortex and 301 000 nm^2^ in the primary motor cortex (Table 7).

The present results show that most synapses presented a macular shape (taking both tissue types together, 83% were macular), whereas 17% were the more complex-shaped synapses (perforated, horseshoe and fragmented), which is comparable to previous reports in other brain areas and species (Geinisman et al. 1987; Jones et al. 1991; Neuman et al. 2016; Hsu et al. 2017; Calì et al. 2018; Santuy et al. 2018a; Domínguez-Álvaro et al. 2019; Montero-Crespo et al. 2020).

Complex-shaped synapses are larger than macular ones, and they have more AMPA and NMDA receptors than macular synapses. Therefore, complex-synapses are thought to constitute a relatively ‘powerful’ population of synapses with more long-lasting memory-related functionality than macular synapses (Geinisman et al. 1987, 1991, 1992a, 1992b, 1993; Lüscher et al. 2000; Toni et al. 2001; Ganeshina et al. 2004a, 2004b; Spruston 2008). Our present results also showed, in both the autopsy and the biopsy samples, that complex-shaped AS were larger than the macular AS, as was previously reported in other brain areas and species (Santuy et al. 2018a; Domínguez-Álvaro et al. 2019; Montero-Crespo et al. 2020).

In addition, perforated synapses were more frequent in the autopsy samples than in the biopsy samples, and they were also more frequent in AS than in SS. As mentioned above, the autopsy cases were older than the biopsy cases. Thus, the frequency of perforated AS in autopsies could be related to both the fixation process and aging, given that advancing age is associated with a lower number of thin spines (which are the main postsynaptic target of macular AS (Benavides-Piccione et al. 2013).

### Postsynaptic targets

A clear preference of glutamatergic axons (forming AS) for spines and GABAergic axons (forming SS) for dendritic shafts was observed, which has also been found in a variety of cortical regions and species (reviewed in DeFelipe et al. 2002). In the present study, the co-analysis of the synaptic type and postsynaptic target showed that the proportions of AS established on spines (‘axospinous’) were around 68% and 78% in the autopsy and biopsy samples, respectively. This difference might be explained by the lower number of spines associated with aging (e.g., see Benavides-Piccione et al. 2013) and/or differences in the preservation of the ultrastructure.

Using the same FIB/SEM technology, in layer II of the human transentorhinal cortex, it was found that 77% of the AS were established on spines (Domínguez-Álvaro et al. 2021). This percentage was lower for the AS in layers II and III of the human entorhinal cortex — 60% and 54%, respectively (Domínguez-Álvaro et al. 2021). These differences and similarities in the proportion of AS on spines and dendritic shafts may represent another microanatomical specialization of the cortical regions (and layers) examined.

Using serial EM, Yakoubi et al. (2018, 2019) analyzed 150 AS in layer IV and 147 AS in layer V of the human temporal neocortex, reporting that axospinous AS account for approximately 79% and 85% of the AS, respectively (Table 8). These results are similar to those found in the frontal cortex of the rhesus monkey (∼80% axospinous AS; Peters et al. 2008; Dumitriu et al. 2010; Hsu et al. 2017) and higher than those found in the striate cortex of the *Macaca mulatta* (64%; Beaulieu et al., 1992). In the present study, we found (70–79% axospinous AS, from autopsy and biopsy brain tissue, respectively, which is a lower percentage than that found in the study of Yakoubi. Since our results come from the analyses of 2128 AS in layer III, the differences between the findings of Yakoubi et al. and the present results could be related to the number of synapses analyzed (Yakoubi et al. analyzed 150 AS in layer IV and 147 AS in layer VI) and/or to differences in the synaptic organization of the layers examined.

Finally, in rodents, this percentage is higher than in primates — for example, in the somatosensory cortex of the young rat (84% axospinous AS; Santuy et al. 2018b) and adult mice (86% axospinous AS; Calì et al. 2018).

Concerning differences between cortical areas, layers and species with regard to the targets of SS, it is more difficult to interpret because of the scarcity of SS. As shown in Table 8, most studies are based on analyses of very few SS (4 to 993 synapses). Our data comes from 176 serially reconstructed SS and we reported 85% of SS established on dendritic shafts. Other studies from our laboratory with similar, or even larger, SS datasets have reported similar axodendritic SS percentages in the human transentorhinal cortex (in which 87% of the 102 SS analyzed were axodendritic SS; Domínguez-Álvaro et al. 2021), and entorhinal cortex (where 86% of the 212 SS analyzed were axodendritic SS; Domínguez-Álvaro et al. 2021). Furthermore, a lower axodendritic SS percentage has been reported in species other than human, including the rhesus monkey (in which 76% of the 354 SS analyzed (in single sections) in the frontal cortex were axodendritic SS; Peters et al. 2008) and the rat (in which 75% of the 574 serially reconstructed SS analyzed in the somatosensory cortex were axodendritic; Santuy et al. 2018b). Thus, it may be that GABAergic synapses are organized differently between humans and the other animal species, with human GABAergic circuits being more focused on axodendritic synapses than in the other animal species.

Finally, the present work constitutes a detailed description of the synaptic organization of the neuropil of layer IIIA of the human BA21. Our study is based on the analysis of a large number of synapses (4945 synapses) that were fully reconstructed in 3D at the ultrastructural level. These results therefore represent robust data for this particular layer. Further studies of the rest of the layers are necessary to further unravel the synaptic organization of this cortical region.

## Ethics approval

Brain tissue samples were obtained following the guidelines and approval of the Institutional Ethical Committees from all involved institutions: School of Medicine, University of Castilla-la Mancha, (Albacete, Spain); and Hospital de la Princesa (Madrid, Spain).

## Availability of data and materials

Most data are available in the main text or the supplementary materials. The datasets used and analyzed during the current study are published in the EBRAINS Knowledge Graph (DOI: 10.25493/5F04-N97):

Dominguez-Alvaro M, Montero M, Alonso-Nanclares L, Blazquez-Llorca L, Rodriguez R, DeFelipe J. (2020). Densities and 3D distributions of synapses using FIB/SEM imaging in the human neocortex (Temporal cortex, T2) [Data set]. Human Brain Project Neuroinformatics Platform.

## Authors’ contributions

JD and LA-N designed and oversaw the project. LA-N designed and carried out the experiments. NC-A performed the data analysis. NC-A and LA-N drafted the initial manuscript. All authors read, reviewed, edited, and approved the final manuscript.

## Acknowledgments

We would like to thank Marta Domínguez-Álvaro for her scientific support, Carmen Álvarez and Lorena Valdés for their technical assistance, and Nick Guthrie for his excellent text editing.

## Abbreviation list

3D: three-dimensional
AS: asymmetric synapses
CA: Cornu Ammonis 1
CF: counting frame
CSR: Complete Spatial Randomness
FIB/SEM: focused ion beam/scanning electron microscopy
KS: Kolmogorov-Smirnov
MW: Mann-Whitney
PB: phosphate buffer
PSD: postsynaptic density
SAS: synaptic apposition surface
SD: standard deviation
SEM: standard error of the mean
TEM: transmission electron microscopy
SS: symmetric synapses.

---

**Supplementary Table 1.** Data from the ultrastructural analysis of neuropil from layer III of Brodmann’s area 21 for individual cases. Data in parentheses are not corrected for shrinkage AS: asymmetric synapses; CF: counting frame; SD: standard deviation; SS: symmetric synapses.

| Case | No. of AS | No. of SS | No. of all synapses | % AS<br>(mean ± SD) | % SS<br>(mean ± SD) | CF<br>volume<br>($\mu\text{m}^3$) | No. of AS/$\mu\text{m}^3$<br>(mean ± SD) | No. of SS/$\mu\text{m}^3$<br>(mean ± SD) | No. of synapses/<br>$\mu\text{m}^3$<br>(mean ± SD) | Distance to<br>the nearest<br>neighbor<br>(nm; mean±<br>SD) |
| --- | --- | --- | --- | --- | --- | --- | --- | --- | --- | --- |
| **AB1** | 756 | 29 | 605 | 95.18±1.63 | 4.82±1.63 | 1146<br>(1032) | 0.51±0.12<br>(0.57±0.13) | 0.03±0.01<br>(0.03±0.01) | 0.54±0.12<br>(0.60±0.12) | 835±58<br>(810±56) |
| **AB2** | 372 | 23 | 395 | 94.13±4.09 | 5.87±4.09 | 794 (715) | 0.47±0.06<br>(0.52±0.07) | 0.03±0.02<br>(0.03±0.02) | 0.50±0.05<br>(0.55±0.05) | 760±86<br>(737±83) |
| **AB3** | 307 | 30 | 337 | 91.04±1.55 | 8.96±1.55 | 764 (688) | 0.40±0.03<br>(0.45±0.03) | 0.04±0.00<br>(0.04±0.01) | 0.44±0.02<br>(0.49±0.03) | 857±94<br>(831±91) |
| **H38** | 614 | 47 | 661 | 92.83±5.19 | 7.17±5.19 | 1016<br>(915) | 0.61±0.18<br>(0.68±0.19) | 0.04±0.02<br>(0.05±0.03) | 0.66±0.17<br>(0.73±0.19) | 707±72<br>(685±70) |
| **H18** | 638 | 46 | 684 | 93.80±3.92 | 6.20±3.92 | 1084<br>(977) | 0.59±0.09<br>(0.65±0.10) | 0.04±0.03<br>(0.05±0.04) | 0.63±0.12<br>(0.70±0.13) | 721±59<br>(710±57) |
| **H60** | 738 | 38 | 776 | 95.18±1.36 | 4.82±1.36 | 1136<br>(1023) | 0.65±0.03<br>(0.72±0.04) | 0.03±0.01<br>(0.04±0.01) | 0.68±0.04<br>(0.76±0.05) | 703±37<br>(682±36) |
| **H47** | 594 | 56 | 650 | 91.48±2.46 | 8.52±2.46 | 1115<br>(1004) | 0.53±0.05<br>(0.59±0.06) | 0.05±0.02<br>(0.06±0.02) | 0.58±0.07<br>(0.65±0.07) | 747±31<br>(725±30) |
| **H94** | 779 | 58 | 837 | 93.01±1.26 | 6.99±1.26 | 1087<br>(979) | 0.72±0.04<br>(0.80±0.05) | 0.05±0.01<br>(0.06±0.01) | 0.77±0.05<br>(0.86±0.05) | 718±14<br>(696±14) |

**Supplementary table 2.** Area of the SAS (nm^2^) in layer III of Brodmann’s area 21 for individual cases. Data in parentheses are not corrected for shrinkage. AS: asymmetric synapses; SAS: synaptic apposition surface; SEM: standard error of the mean; SS: symmetric synapses.

| Case | Type of synapse | Area of SAS<br>(mean ± SEM) |
| --- | --- | --- |
| AB1 | AS | 85 257±2392<br>(79 289±2224) |
| | SS | 37 829±3674<br>(35 181±3417) |
| AB2 | AS | 116 131±4473<br>(108 002±4160) |
| | SS | 57193±9384<br>(53 190±8728) |
| AB3 | AS | 132 104±5626<br>(122 857±5232) |
| | SS | 84 027±10474<br>(78 145±9741) |
| H38 | AS | 113 952±4266<br>(10 5976±3967) |
| | SS | 82 397±6765<br>(76 629±6292) |
| H18 | AS | 90 727±3161<br>(84 376±2940) |
| | SS | 53 326±5085<br>(49 594±4729) |
| H60 | AS | 106 981±1509<br>(99 492±1403) |
| | SS | 78 535±7711<br>(73 038±7171) |
| H47 | AS | 130 042±1680<br>(120 939±1563) |
| | SS | 93 887±7308<br>(87 315±6797) |
| H94 | AS | 106 746±3348<br>(99 274±3114) |
| | SS | 98 375±7874<br>(91 489±7323) |

**Supplementary table 3.**
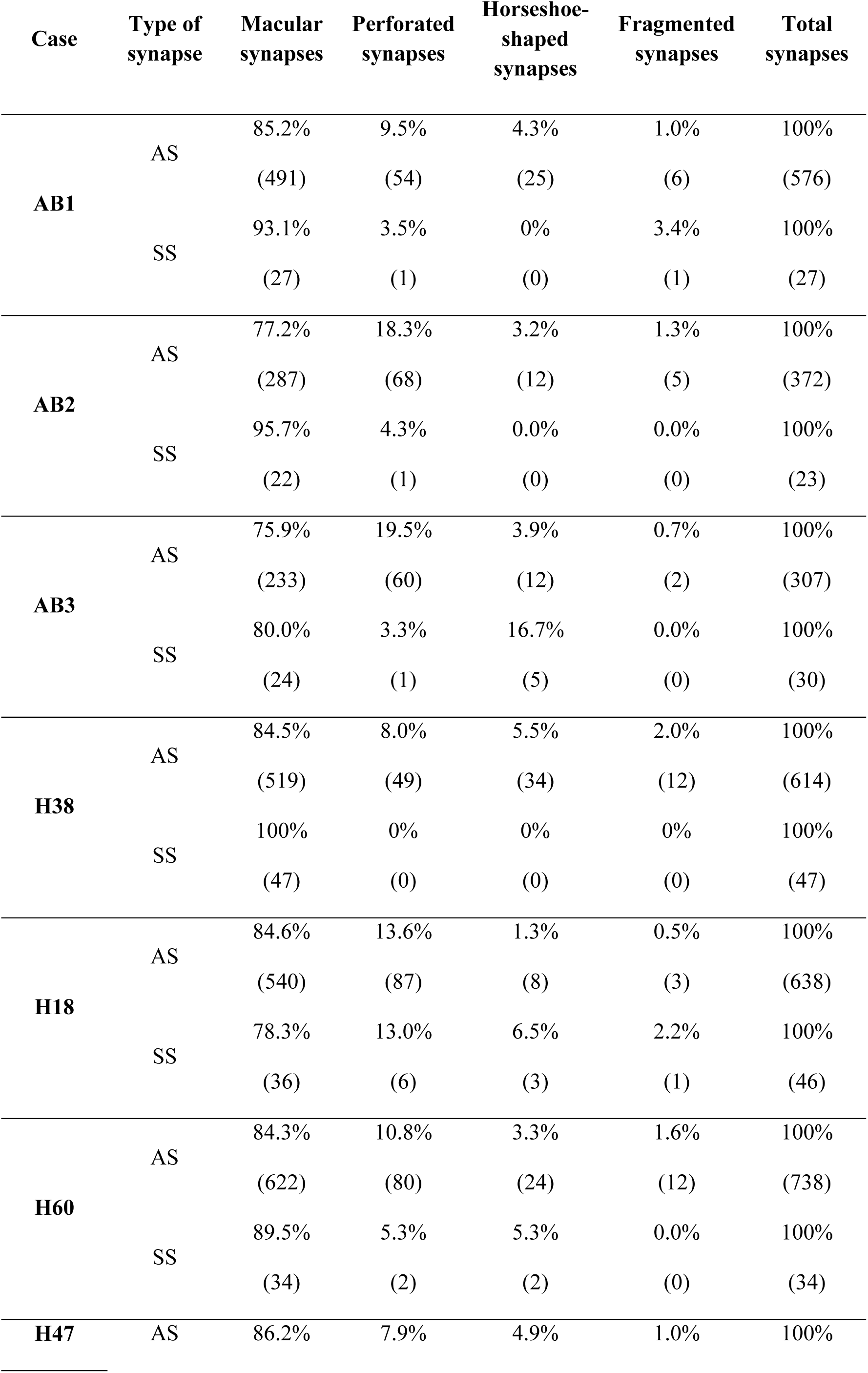
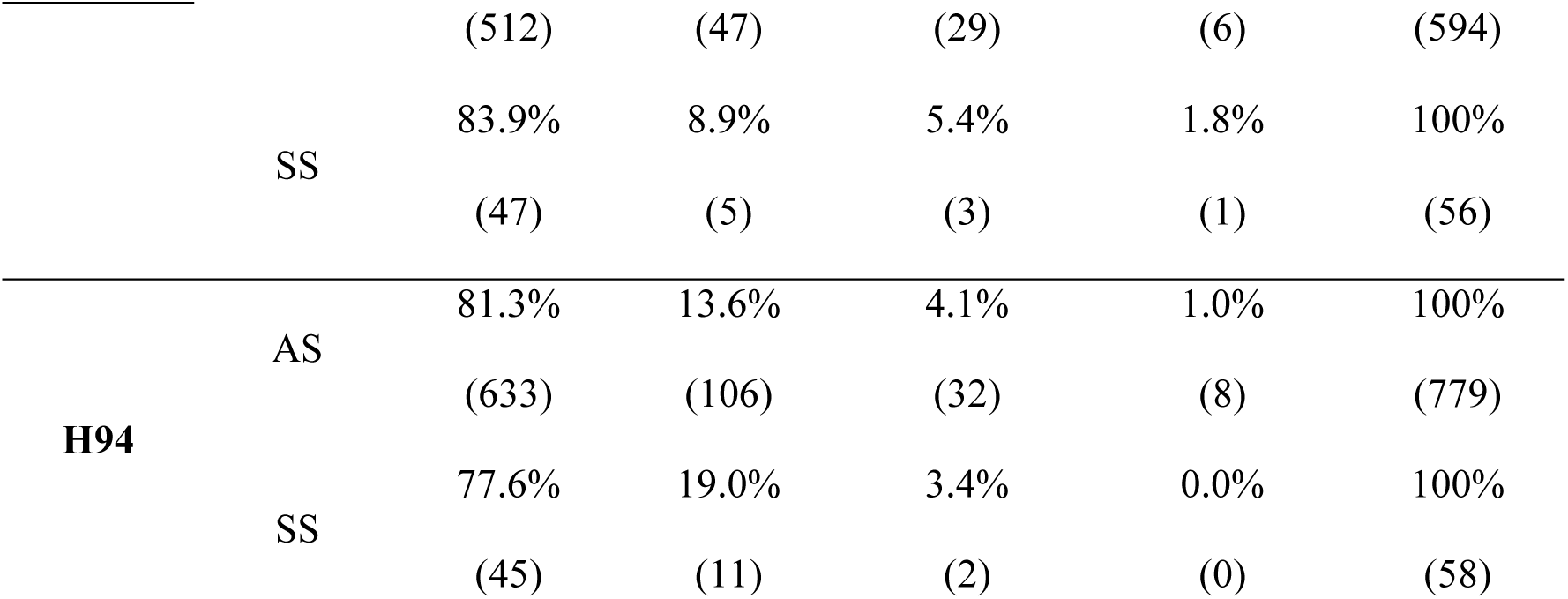
Proportion of macular, perforated, horseshoe-shaped and fragmented synapses in layer III of Brodmann’s area 21 for individual cases. Data are given as percentages with the absolute number of synapses studied in parentheses. AS: asymmetric synapses; SS: symmetric synapses.

**Supplementary table 4.** Distribution of asymmetric (AS) and symmetric (SS) synapses on spines and dendritic shafts in layer III of Brodmann’s area 21 for individual cases. Synapses on spines have been sub-divided into those established on spine heads that are fully reconstructed or non-fully reconstructed and those that are established on spine necks. Moreover, we differentiated between aspiny and spiny dendritic shafts. Data are expressed as percentages with the absolute number of synapses studied given in parentheses. AS: asymmetric synapses; SS: symmetric synapses.

| Case | Type of synapse | Synapses on spine heads | Synapses on putative spine heads | Synapses on spine necks | Synapses on dendritic shaft with spines | Synapses on dendritic shaft without spines | Total synapses |
| --- | --- | --- | --- | --- | --- | --- | --- |
| **AB1** | AS | 53.2% | 12.9% | 0.9% | 20.8% | 12.3% | 100% |
|  |  | (182) | (44) | (3) | (71) | (42) | (342) |
|  | SS | 15.0% | 0.0% | 0.0% | 75.0% | 10.0% | 100% |
|  |  | (3) | (0) | (0) | (15) | (2) | (20) |
| **AB2** | AS | 47.3% | 23.8% | 0.9% | 11.9% | 16.1% | 100% |
|  |  | (159) | (80) | (3) | (40) | (54) | (336) |
|  | SS | 4.3% | 4.3% | 4.3% | 52.2% | 34.8% | 100% |
|  |  | (1) | (1) | (1) | (12) | (8) | (23) |
| **AB3** | AS | 51.2% | 16.0% | 2.5% | 23.1% | 7.1% | 100% |
|  |  | (144) | (45) | (7) | (65) | (20) | (281) |
|  | SS | 19.2% | 3.8% | 3.8% | 42.3% | 30.8% | 100% |
|  |  | (5) | (1) | (1) | (11) | (8) | (26) |
| **H38** | AS | 68.3% | 24.5% | 0.0% | 2.9% | 4.3% | 100% |
|  |  | (95) | (34) | (0) | (4) | (6) | (139) |
|  | SS | 0.0% | 10.0% | 5.0% | 40.0% | 45.0% | 100% |
|  |  | (0) | (2) | (1) | (8) | (9) | (20) |
| **H18** | AS | 47.5% | 19.8% | 1.8% | 12.9% | 18.0% | 100% |
|  |  | (103) | (43) | (4) | (28) | (39) | (217) |
|  | SS | 6.9% | 0.0% | 0.0% | 20.7% | 72.4% | 100% |
|  |  | (2) | (0) | (0) | (6) | (21) | (29) |
| **H60** | AS | 58.0% | 25.1% | 1.0% | 5.8% | 10.1% | 100% |
|  |  | (120) | (52) | (2) | (12) | (21) | (207) |
|  | SS | 0.0% | 11.1% | 0.0% | 44.4% | 44.4% | 100% |
|  |  | (0) | (1) | (0) | (4) | (4) | (9) |
| **H47** | AS | 50.0% | 20.3% | 0.0% | 13.9% | 15.8% | 100% |
|  |  | (79) | (32) | (0) | (22) | (25) | (158) |
|  | SS | 0.0% | 0.0% | 8.3% | 33.3% | 58.3% | 100% |
|  |  | (0) | (0) | (1) | (4) | (7) | (12) |
| **H94** | AS | 58.7% | 20.8% | 0.4% | 9.2% | 10.9% | 100% |
|  |  | (263) | (93) | (2) | (41) | (49) | (448) |
|  | SS | 8.1% | 10.8% | 0.0% | 21.6% | 59.5% | 100% |
|  |  | (3) | (4) | (0) | (8) | (22) | (37) |

**Supplementary Figure 1:**
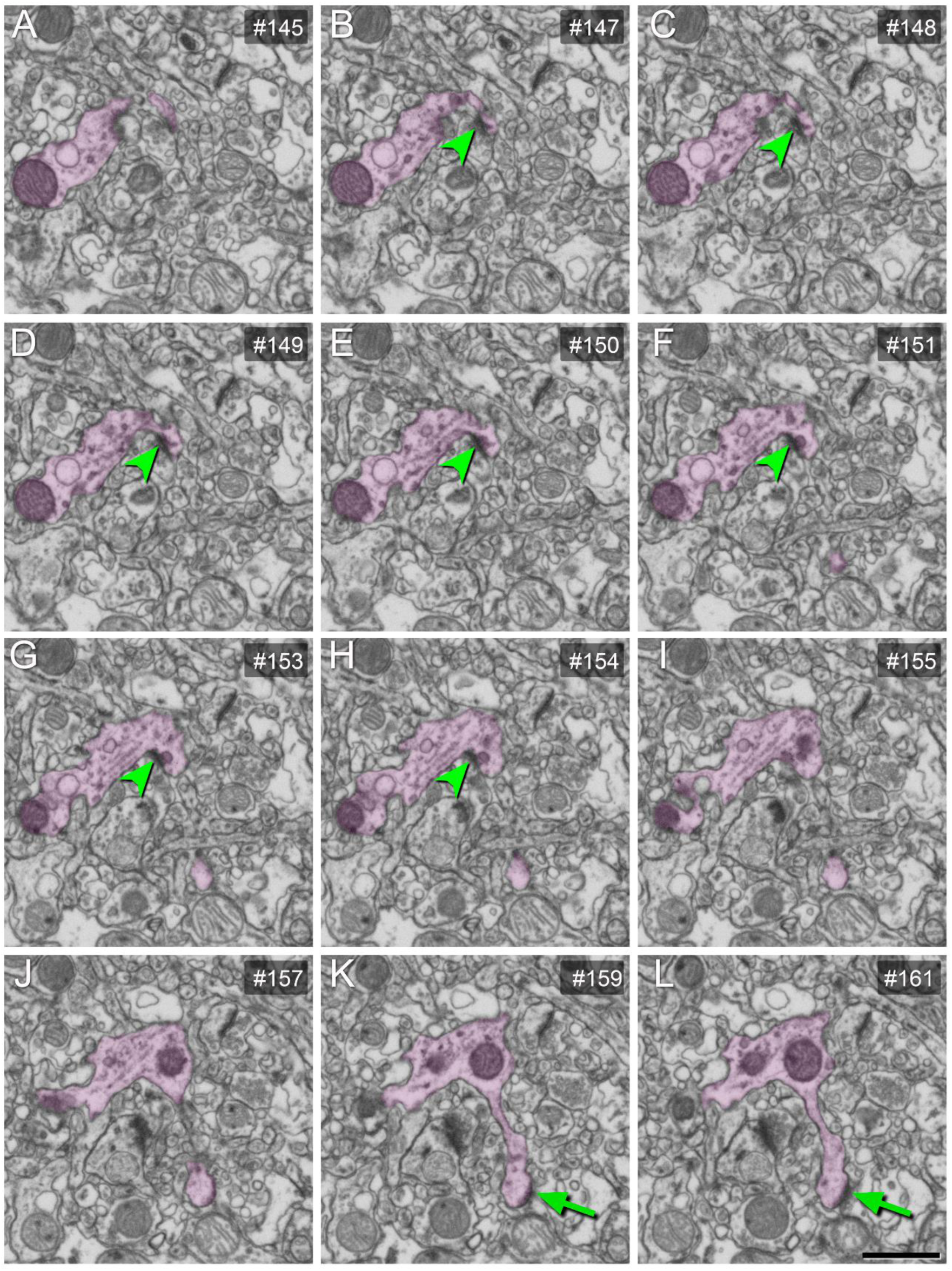
serial images obtained by FIB/SEM showing a dendritic segment with a dendritic spine (clear pink). AS established on a spine neck (arrowhead) and on a spine head (arrow) are illustrated. AS: asymmetric synapse. Scale bar (in L) indicates 1000 nm in A–L.

**Supplementary Figure 2:**
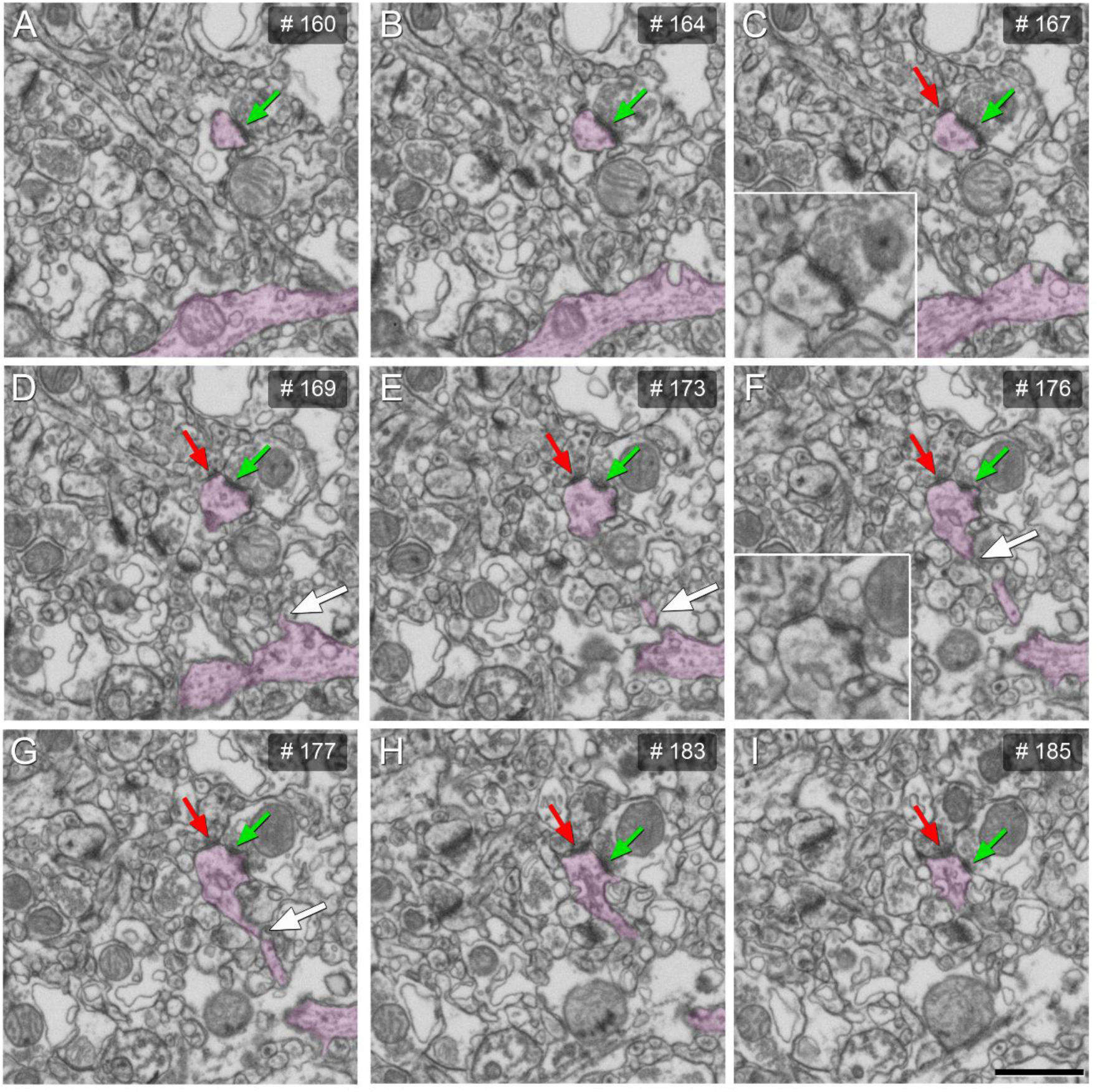
serial images obtained by FIB/SEM showing a dendritic spine head emerging from its parent dendrite (clear pink). AS (green arrow) and SS (red arrow) are established on the same spine head. The white arrow indicates the dendritic spine neck. AS, asymmetric synapse; SS, asymmetric synapse. Scale bar (in I) indicates 1000 nm in A–I, and 1600 nm in the expanded frame of C and F.

**Supplementary Figure 3.**
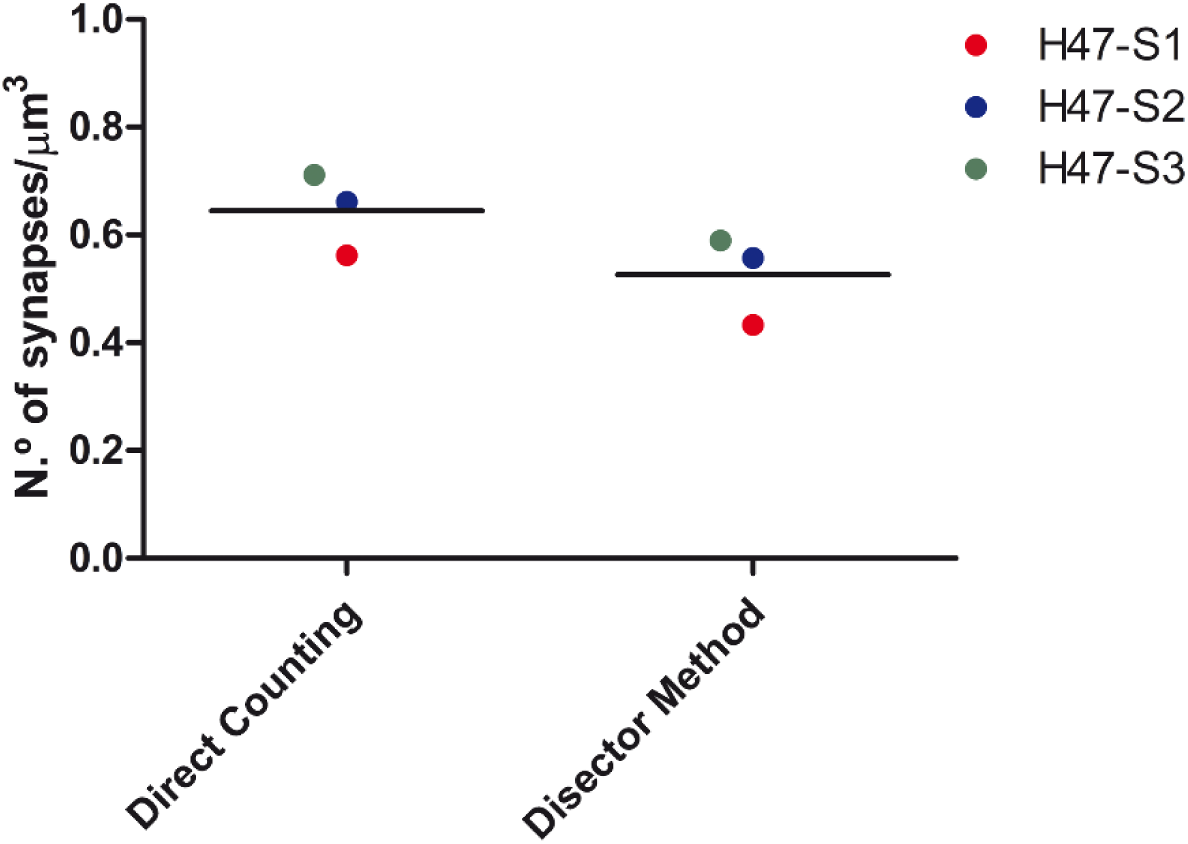
Graph showing comparison of the mean synaptic density obtained by direct counting of synapses and by disector method. All data were calculated for the same stacks of images (S1, S2 and S3) from a single biopsy case (H47). Direct counting of synapses was carried out within a counting frame for 3 stacks of serial images. The disector method obtains the estimations by means of 101 pairs of disectors whose h-value was 3 times the section thickness (60nm). Data obtained from the direct counting of the synapses are unique for each tissue brick composed by the stack of images, while data obtained by the disector method are derived from multiple estimations. Values are not corrected for shrinkage to facilitate comparisons with the literature.

**Supplementary Figure 4.**
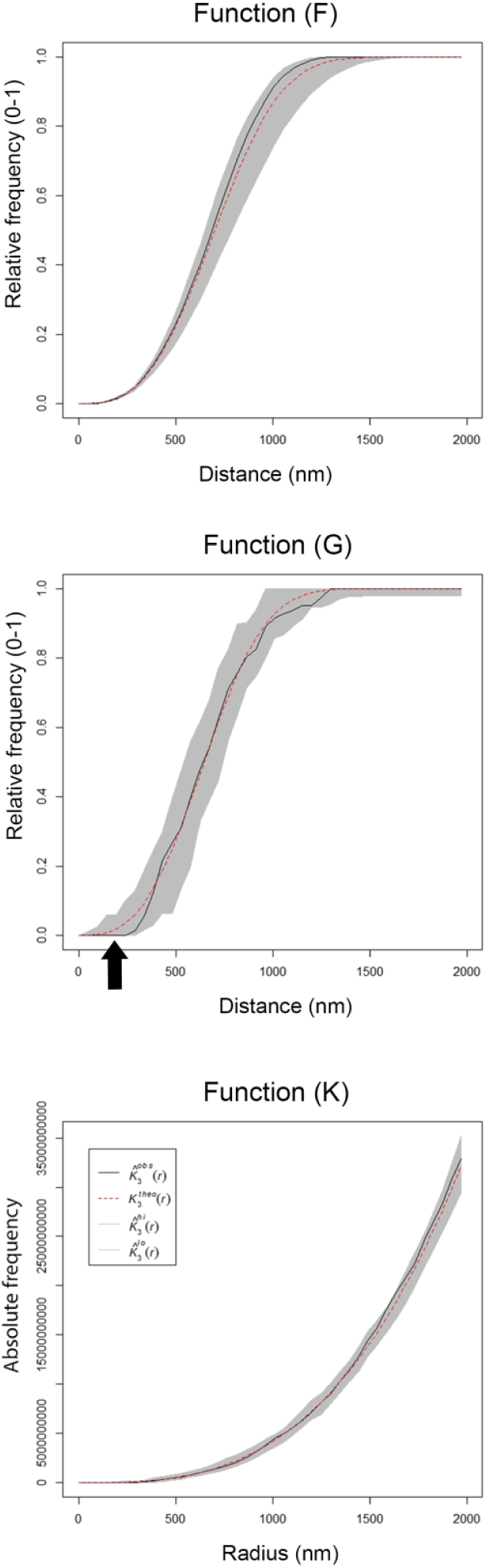
Analysis of the 3D synaptic spatial distribution in layer III of Brodmann’s area 21. Red dashed traces correspond to a theoretical homogeneous Poisson process for each function (F, G, K). The black continuous traces correspond to the experimentally observed function in the sample. The shaded areas represent the envelopes of values calculated from a set of 99 simulations. All plots show a distribution which fit into a Poisson function. In the G function, the arrow shows a dead space due to the fact that synapses cannot be too close to each other since they cannot overlap in space. All data were obtained from case AB2.

**Supplementary Figure 5.**
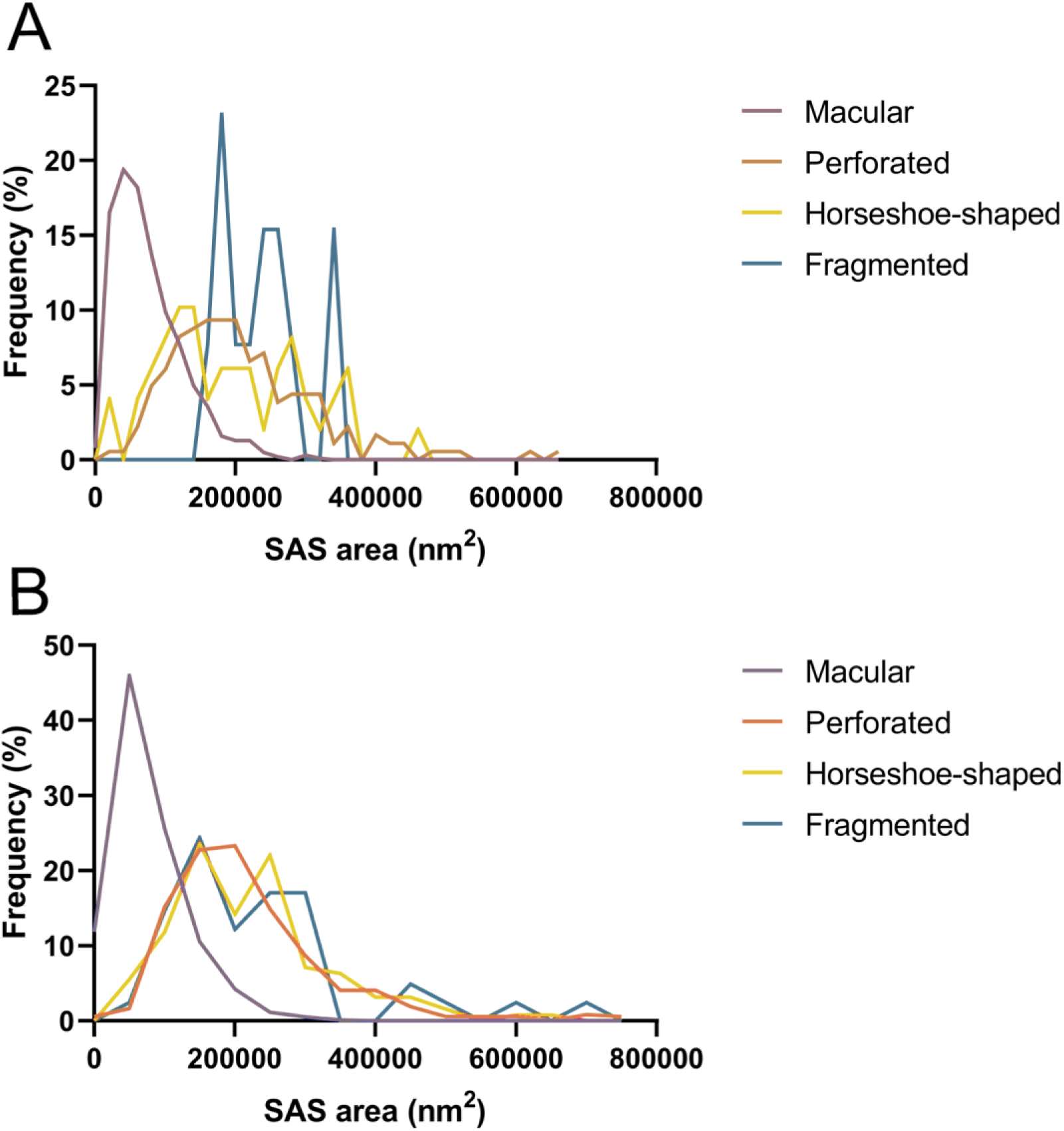
Frequency histograms of the SAS area of macular, perforated, horseshoe-shaped and fragmented AS from autopsy (A) and biopsy (B) samples, in layer III of Brodmann’s area 21. The AS SAS area of the macular synapses were significantly smaller than in the case of perforated, horseshoe-shaped and fragmented synapses, in both autopsy samples (KW, p<0.0001) and biopsy samples (KW, p<0.0001). AS: asymmetric synapses; SAS: synaptic apposition surface.

**Supplementary Figure 6.**
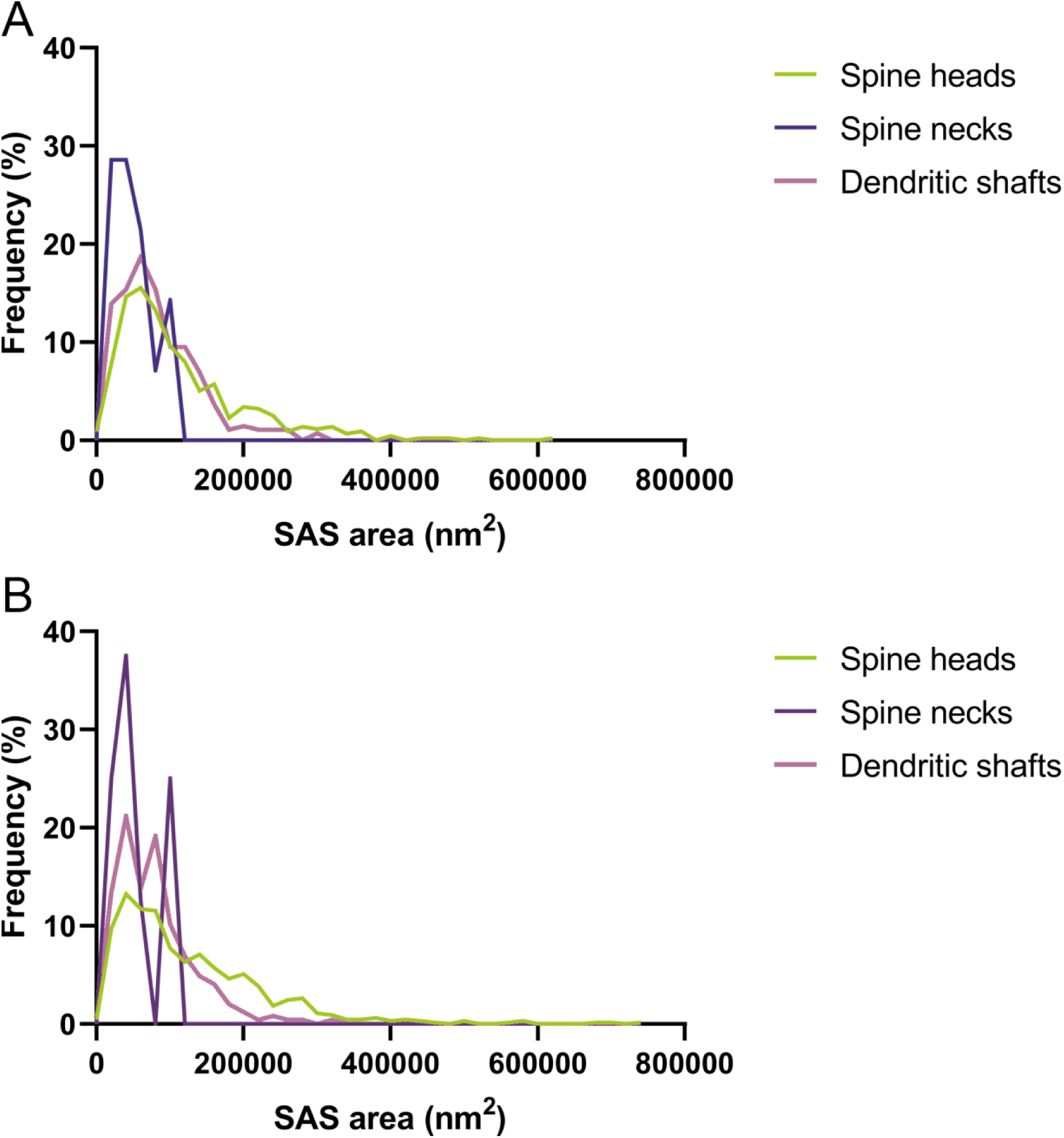
Frequency histograms of the SAS area of AS targeting spine heads, spine necks and dendritic shafts from autopsy (A) and biopsy (B) samples, in layer III of Brodmann’s area 21. The SAS area of the AS established on spine heads were significantly larger (KW, p<0.05) than AS established on spine necks or dendritic shafts for both the autopsy and the biopsy samples. AS: asymmetric synapse; SAS: synaptic apposition surface.

